# Extracellular matrix stiffness regulates degradation of the Hippo kinase MST2 via SCF ^βTrCP^

**DOI:** 10.1101/2021.01.31.429078

**Authors:** Ana Paula Zen Petisco Fiore, Ana Maria Rodrigues da Silva, Helder Veras Ribeiro Filho, Antonio Carlos Manucci, Pedro de Freitas Ribeiro, Mayara Carolinne Silva Botelho, Paulo Sergio Lopes de Oliveira, Michele Pagano, Alexandre Bruni-Cardoso

## Abstract

Tumor microenvironments display disrupted mechanical properties, including altered extracellular matrix (ECM) rigidity. ECM stiffening perturbs cell tensional homeostasis resulting in activation of mechanosensing transcriptional co-activators, such as the Hippo pathway effectors YAP and TAZ. The Hippo pathway plays central roles in development and tumorigenesis, but how the proteostasis of the Hippo kinase MST2 is regulated remains unknown. Here, we show that ECM stiffness induces MST2 degradation via proteasome degradation. MST2 degradation is enhanced in human breast epithelial cells (HMEC) that are cultured in stiffer microenvironments due to integrin and integrin-linked kinase activation. MST2 knockdown resulted in increased nucleus-to-cytoplasm ratio of YAP in physiological and breast tumor rigidities and altered mechanoregulated cellular processes in HMEC. We found that MST2 is ubiquitinated by the SCF^βTrCP^ ubiquitin ligase. Site-directed mutagenesis combined with computational molecular dynamics studies revealed that βTrCP binds MST2 via a noncanonical degradation motif. Our study uncovers the underlying biochemical mechanisms controlling MST2 degradation and demonstrates how changes in the microenvironment rigidity regulate the proteostasis of a central Hippo pathway component.

## Introduction

The tissue microenvironment is a source of mechanical signals, such as those provided by stiffness of the extracellular matrix (ECM), that regulate cell shape and several cellular processes (*1, 2*). Cells sense composition and mechanical properties of the ECM eliciting adjustment in tension and rearranging the actomyosin cytoskeleton (*3, 4*). Altered ECM mechanical properties are found in many diseases, including cancers (*5–7*). As cancer progresses, malignant cells disturb the surrounding tissue physical properties (*5*). Reciprocally, mounting evidence has shown that tissue stiffening can induce phenotypic transformation (*4, 5, 8, 9*). Commonly used as diagnosis and prognosis of several types of cancer, increased tissue stiffness is caused by ECM deposition and remodeling. And, a stiff ECM activates nuclear mechanosensing molecular relays, such as the co-activator factors YAP (Yes-associated protein)/TAZ (transcriptional co-activator with PDZ-binding motif, also known as WWTR1) (*10*), which promote proliferation and invasion (*5, 8, 9*).

In epithelial cells exposed to physiological rigidities, YAP/TAZ are predominantly cytoplasmic but translocate to the nucleus in stiff microenvironments (*11*). YAP/TAZ subcellular localization and transcriptional activity are modulated by Hippo, a pathway cored by the kinases MST1/2 and LATS1/2. On the other hand, YAP/TAZ mechanosignaling promotes fibrosis, resulting in microenvironment stiffening (*12–24*). The transduction of mechanical signals that results in cell shape remodeling and in changes in gene expression are initially sensed by the ECM receptors, integrins, which in turn trigger assembly of focal adhesions (FAs), multiprotein complexes linked to filamentous actin (*2, 25*). FAs are composed of Focal Adhesion Kinase (FAK) and/or Integrin-Linked Kinase (ILK), integrin clusters and scaffold proteins, and are modulated by substrate rigidity in response to intracellular tension built in the actomyosin cytoskeleton (*26, 27*). FAK and ILK phosphorylate multiple substrates and help converge integrin and growth factor-receptor triggered signaling pathways (*28*).

Although a great deal is known regarding regulation of YAP/TAZ levels and activities by the ECM stiffness, little is known about the mechanical regulation of the Hippo pathway components. We sought at investigating whether MST2, a core Hippo serine-threonine kinase, was modulated by microenvironment rigidity and found that stiff microenvironments induces MST2 proteolysis in human mammary epithelial cells (HMEC). Silencing of *STK3*, the gene encoding MST2, resulted in increased nucleus/cytoplasm ratio of YAP in physiological (Low) and breast-tumor rigidities (Mid), increased proliferation rates in all tested degrees of stiffness, changes in cell size and shape, and increased F-actin alignment in HMEC cultured in intermediate stiffness. We found that MST2 is ubiquitinated by the E3-ligase complex Skp1/Cul1/F-box-protein/β-transducing-repeat-containing protein (SCF^βTrCP^) and degraded via the proteasome. The E3-ligase substrate-recognition subunit βTrCP binds to MST2 via a noncanonical degradation motif (degron). We also found that signaling from the integrin-ILK (integrin-linked kinase) axis results in increased MST2 and βTrCP binding, leading to MST2 degradation. Our findings shed light on the mechanisms controlling Hippo activity by targeted proteolysis and discovered that degradation of MST2 is a crucial part in the mechanoregulatory circuitry that senses and transduces physical cues from the cell microenvironment.

## Results

### Stiff microenvironments reduce MST2 levels

We asked whether the level of MST2 was regulated by ECM stiffness. For this, we cultured MCF10A cells, a non-tumoral human mammary epithelial cell (HMCE) line highly sensitive to modulation of substrate stiffness (*8*), on fibronectin-coated polyacrylamide hydrogels that can be tuned in rigidity by changing the concentration of polyacrylamide (*29*). Three degrees of elastic modulus were used: 0.48 KPa (Low; a physiological rigidity), 4.47 KPa (Mid; average rigidity of the mammary tumor stroma) and 40.40 KPa (Hi; a supraphysiological stiffness).

Immunoblotting of MCF10A-cell lysates revealed that levels of MST2 decrease in response to stiffer substrates (Fig. 1A). But, intriguingly, *STK3* (the gene encoding MST2) mRNA levels did not change in response to different rigidities (Fig. 1B), suggesting that regulation of MST2 protein levels could be via proteolytic degradation in different mechanical contexts. We silenced *STK3* using siRNA to evaluate whether modulation of MST2 levels would reflect in alterations in the nuclear to cytoplasm ratio of YAP and also phenotypical changes in MCF10A cells (Fig. 1C, Supp. Fig. 1A and Fig. 1D). In unperturbed MCF10A cells cultured in Low stiffness, YAP is predominantly cytoplasmic, as determined by measuring nuclear to cytoplasmic YAP ratio by immunofluorescence followed by image quantificaion. YAP preferentially localizes in the nucleus in cells grown in stiffer substrates (Fig. 1D-E and Supp. Fig. 1B) (*10*). Silencing *STK3* results in increased nucleus to cytoplasm YAP ratios in Low and Mid stiffness in comparison to cells treated with a control siRNA (siControl). Cell proliferation, as measured by EdU incorporation, increases in Mid and Hi rigidities in comparison to cells cultured in a soft ECM (Fig. 1F and Supp. Fig. 1C). However, silencing *STK3* resulted in higher percentages of EdU-positive cells in all three degrees of rigidity.

**Figure 1:**
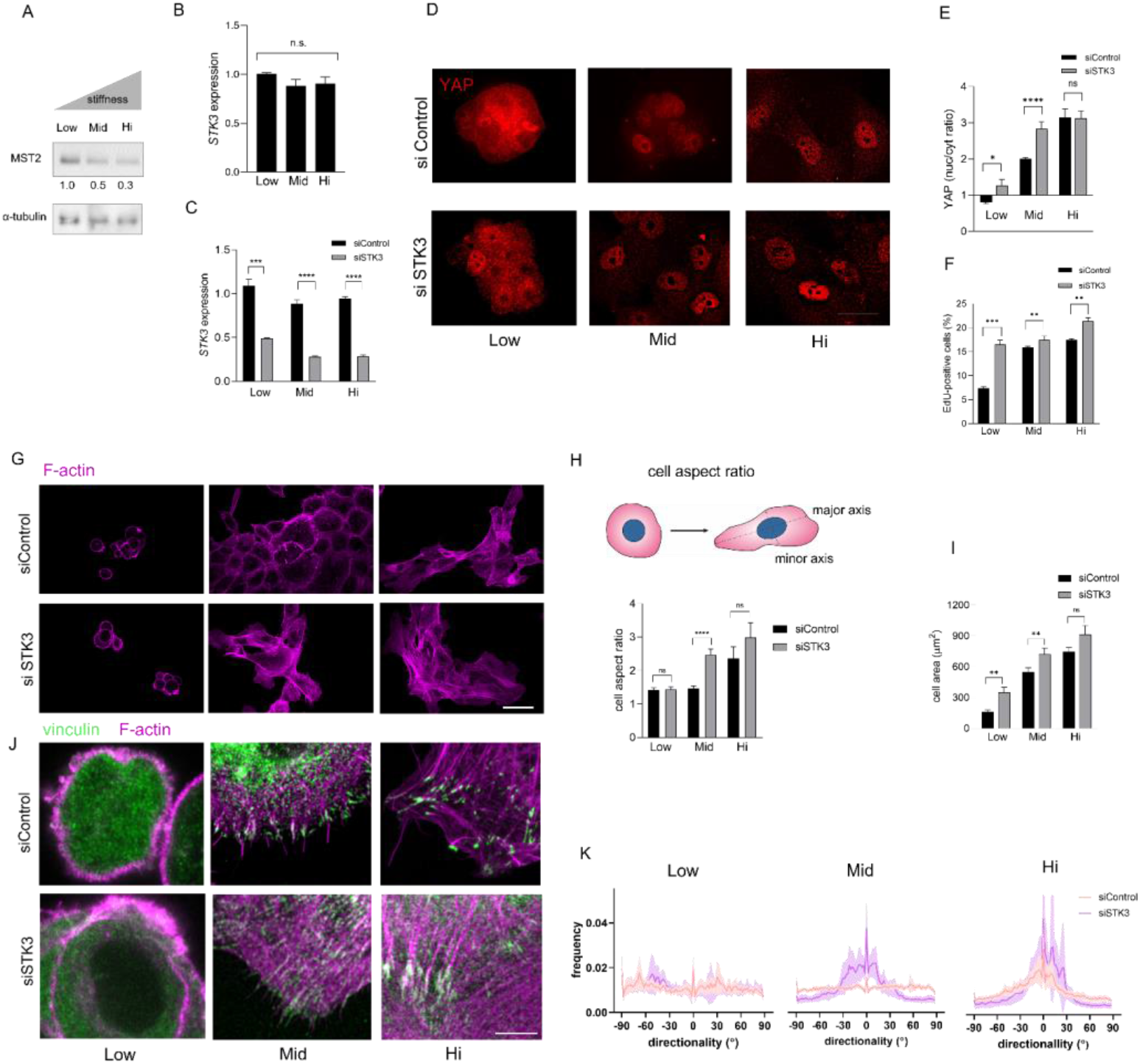
The extracellular matrix stiffness regulates the level of MST2, and *STK3* knockdown alter cellular mechanoregulatory outputs. A) Immunoblots for MST2 and α-tubulin (used as a loading control) of MCF10A cells cultured on fibronectin-1 (FN1) coated hydrogels with different rigidities: 0.48 KPa (Low), 4.47 KPa (Mid) and 40.40 (Hi). Fold changes of MST2 levels are shown under between the immunoblots. B) Quantitative RT-PCR for *STK3* (gene that encodes MST2) of MCF10A cells cultured on hydrogels with different degrees of rigidity. GAPDH was used as an endogenous control to normalize gene expression. Data are expressed as mean ± SEM of e^−ΔΔct^ of (N = 3). C) Quantitative RT-PCR for *STK3* of MCF10A cells silenced for *STK3* cultured on hydrogels. GAPDH was used as an endogenous control to normalize gene expression. MCF10A cells were transfected with an siRNA for *STK3* (siSTK3) or nontargeting siRNA (siControl). Data are expressed as mean ± SEM of e^−ΔΔct^ (N = 3). See also Figure S1A for MST2 protein expression after siRNA treatments. D) Representative microscope images of immunofluorescence (IF) for YAP (red) of MCF10A cells silenced for *STK3* cultured on hydrogels. See Figure 1S2 for images of YAP staining paired with DNA labeling with DAPI. E) Quantification of nuclear-to-cytoplasmic ratio of YAP staining of MCF10A cells silenced for *STK3* (N= 10-15 per condition in each of the 3 independent biological replicates). F) Quantification of EdU-positivity reflecting proliferation of MCF10A cells silenced for *STK3*. (N= 15 40x microscope fields per condition. Each microscope field contained 20-70 cells. The data in the graph are a compilation from 2 independent biological replicates. G) Representative low magnification Airy-Scan superresolution microscope images of MCF10 cells silenced for *STK3* and stained with fluorescent phalloidin for F-actin visualization (magenta). H) Aspect ratio changes of MCF10A cells silenced for *STK3*. Top: schematic depicting a rounded cell and an elliptical cell with highlighting its major and minor axes. Bottom: Quantification of cell aspect ratio (N= 10-15 cells per condition in each of the 3 independent biological replicates). I) Quantification of the area of MCF10A cells silenced for *STK3*. (N= 10-15 cells per condition in each of the 3 independent biological replicates). J) Representative high magnification Airy-Scan superresolution microscope images of MCF10 cells silenced for *STK3* and stained with phalloidin (magenta) and vinculin (green). K) Quantification of F-actin alignment in MCF10A cells silenced for *STK3*. Data are plotted for the three degrees of stiffness as frequency of fibers as a function of their directionality in degrees (°) with respect to the major cell axis (a directionality of 0° means perfect alignment with the major cell axis). (N= 10-15 cells per condition in each of the 3 independent biological replicates). Statistical significance = *p<0.05, **p<0.01, ***p<0.001, ****p<0.0001 and n.s denotes no statistical significance. For data in A, B and F, the comparison applied was ANOVA followed by Dunnett test, while for E, H and I the statistical analysis used was the Kruskal-Wallis test. Scale bars, D =10 μm, G = 40 μm, J = 5 μm.

Knocking down MST2 resulted in striking changes in cell shape particularly in cells cultured in Mid and Hi stiffnesses (Fig. 1G-J). In low stiffness, siControl MCF10A cells behave as a normal epithelium: cells acquire a rounded morphology, F-actin is largely cortical, vinculin (a focal adhesion protein) staining is not defined in a classical focal adhesion (FA) pattern (Fig. 1G and Fig. 1J). On the other hand, in stiffer hydrogels, MCF10A cells are flatter and more spread - despite not losing cell-to-cell contact – display F-actin bundles and a classical focal adhesion pattern of vinculin staining. In Mid and Hi substrates, cells are less rounded as evidenced by increased cell aspect ratio. (Fig. 1G and Fig. 1H). MCF10A cells treated with siSTK3 were larger in Low and Mid rigidities and displayed a less rounded morphology in Mid and Hi in comparison to cells treated with a control siRNA (Fig. 1I). In correspondence to the morphological changes, siSTK3-treated cells displayed in Mid and Hi rigidities pronounced stress fibers that were more aligned (in relation to the cell’s major axes) and colocalized with typical FAs, as revealed by Airy-Scan superresolution microscopy (Fig. 1J-K). Altogether, these data revealed ECM rigidity modulates MST2 levels and that this can affect changes in YAP subcellular localization and result in actin cytoskeleton and morphological changes.

### MST2 is degraded via the proteasome after ubiquitination by SCF^βTrCP^

Because MST2 protein levels decreased in response to stiffer ECMs - albeit its coding mRNA remained unaltered in the same conditions- we decided to investigate whether inhibition of the proteasome with MG132 or cullin-1 neddylation (covalent conjugation of the ubiquitin-like protein NEDD8 required for activation of SCF E3-ligase complexes) with MLN4924 would result in accumulation of MST2 in a panel of non-malignant and malignant HMEC lines. Accordingly, treatment with MG132 or MLN4924 resulted in MST2 accumulation in 4 of 7 human breast cell lines grown on regular cell culture polystyrene plates, which have a rigidity of ~10^9^ P (Fig. 2A).

**Figure 2:**
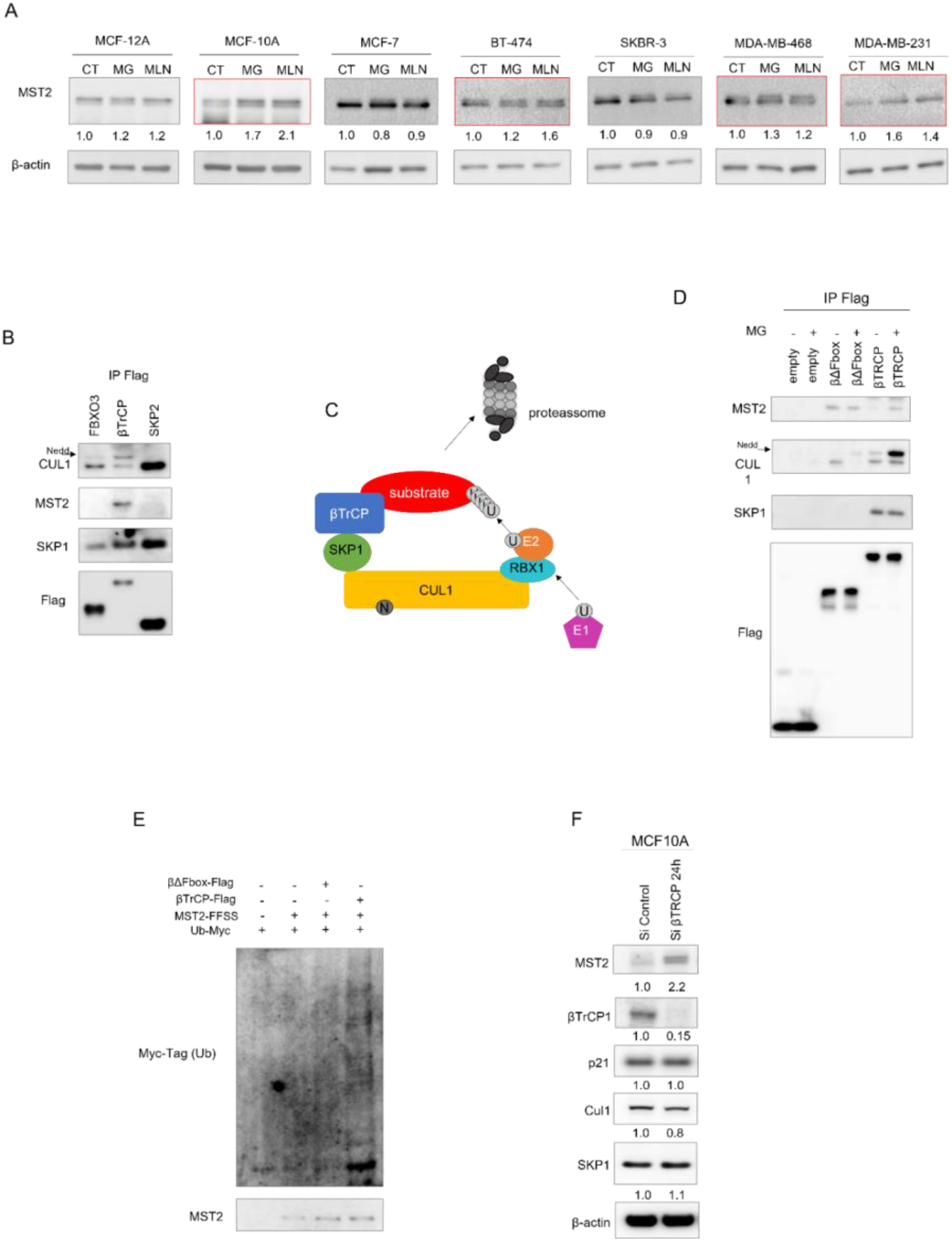
MST2 is ubiquitinated by the SCF^βTrCP^ E3 ligase complex and degraded via proteasome: A) Immunoblots for MST2 in protein extract from breast mammary cell lines cultured in the absence (−) or presence (+) of the proteasome inhibitor MG132 (MG) or the CUL1 neddylation inhibitor MLN4924 (MN) for 3 hours. Cell lines in which MST2 was accumulated upon treatments are highlighted (red outline). β-actin was used as a loading control. B) Co-immunoprecipitation (co-IP) of FBXO3, βTrCP and SKP2 followed by immunoblotting for CUL1, MST2, SKP1 and Flag. HEK293T cells were transfected with Flag-tagged constructs FBXO3, βTrCP and SKP2. Transfected cells were treated for 1.5 hour with MG132, lysed and the immunoprecipitation was performed using M2 antibody coated agarose beads. Both CUL1 and SKP1 are proteins that interact with F-box proteins and were used as Co-IP controls. Flag was detected to assess the success of precipitation of the constructs. C) Schematic of the E3 ligase complex SCF^βTrCP^. As other SKP1-CUL1-Fbox (SCF) complexes, SCF^βTrCP^ consists of 4 main subunits: SKP1, CUL1, RBX1 and the Fbox protein βTrCP that functions as a receptor for target proteins, giving specificity to the complex. Protein degradation by the 26S proteasome depends on a cascade of enzymes. The E1 enzyme binds and activate ubiquitin (U), followed by the recruitment of an E2 ubiquitin-conjugating enzyme that is then loaded with activated ubiquitin. E2 interacts with the E3 complex via the adaptor protein (RBX). After a sequence of ubiquitin conjugation cycles the target protein is degraded by the proteasome. CUL neddylation (N) is required for SCF activation. D) Co-IP confirming the direct binding of MST2 and βTrCP. HEK293T cells were transfected with Flag-tag fused to full length βTrCP or the dominant negative βTrCP (βΔFbox) constructs, were treated or not with MG132 for 1.5 hour. Flag immunoprecipitation was performed using M2 antibody coated beads. Flag and endogenous MST2, CUL1 and SKP1 were detected by immunoblotting. E) Detection of MST2 ubiquitination. HEK293T cells were transfected with Flag-Flag-Strep-Step-tagged MST2 (FFSS-MST2) Flag-tagged β-TrCP or Flag-tagged βΔFbox and Myc-tagged Ubiquitin (Ub-Myc). After a stringent extraction, the FFSS_MST2 was precipitated with tactin-coated beads. To evaluate ubiquitination of MST2, Ub-Myc was detected with immunoblotting using an antibody against Myc-tag. MST2 was also detected by immunoblotting. F) Silencing of βTrCP in MCF10A cells results in MST2 accumulation. Cells were cultured in the presence of siRNA for βTrCP or a non-targeting siRNA (siControl). Immunoblots confirming that silencing βTrCP was efficient and resulted in increased MST2 levels. Immunoblot of p21 was used as a negative control, since it is not a βTrCP substrate. SKP1, p21 and CUL1 did not vary with βTrCP silencing. β-actin was used as loading control. A and F) Fold changes of protein levels are shown under their respective immunoblots. All cell culture assays in this figure were performed on regular cell culture polystyrene plates and flasks.

To identify which specific E3-ligase complex would be responsible for ubiquitinating MST2, we assessed the literature and available MST2 interactomes. Mass spectrometry analysis performed in our laboratory and by others independently identified three E3-ligase substrate-recognition subunits in the MST2 interactomes: FBXO3, SKP2 and βTrCP (*30–32*). Furthermore, a recent report demonstrated that βTrCP binds MST2 in an precipitation assay of ectopically expressed proteins and that inhibition of the proteasome resulted in MST2 accumulation in HEK293T cells (*33*). We transfected HEK293T cells with flag-tagged βTrCP, FBXO3, or SKP2 constructs and performed immunoprecipitation assays followed by immunoblotting (Fig. 2B). Endogenous MST2 was detected only in the flag-tagged βTrCP immunoprecipitants. βTrCP is part of the SCF^βTrCP^ E3-ligase complex which tags key regulatory proteins of proliferation (including YAP/TAZ (*34*)), DNA repair and apoptosis for degradation (*35–37*). As other SCF complexes, SCF^βTrCP^ consists of 4 main subunits: SKP1, CUL1, RBX1 and the Fbox protein βTrCP, the latter acts as a substrate receptor, providing specificity to the complex (*38*) (Fig. 2C). To confirm the direct interaction of βTrCP and MST2, we transfected HEK293T cells with a full length βTrCP or a βTrCP lacking the whole Fbox domain (βΔFbox), a sequence of ~40 amino acids that is required for Fbox proteins to bind SKP1 and, hence, to assemble the SCF complex (*39*). Co-IP followed by immunoblotting revealed that MST2 binds to βΔFbox showing a more intense band than the βTrCP lane (Fig. 2D). Proteasome inhibition increased the interaction between βTrCP and MST2, but resulted in no increment of βΔFbox and MST2 binding. This occurred, probably, because substrates bound to βΔFbox are not ubiquitinated. These co-IP experiments confirmed that MST2 interacts with βTrCP even in the absence of the intact SCF^βTrCP^ complex, indicating that MST2 binds directly βTrCP and not via other SCF subunits. A ubiquitination assay using Myc-tagged ubiquitin confirmed that MST2 is, indeed, ubiquitinated only in the presence of βTrCP but not βΔFbox (Fig. 2E).

Next, we assessed if our findings of βTrCP binding and regulation of MST2 levels in HEK293T cells would be consistent in a HMEC without ectopic overexpression. Silencing βTrCP in MCF10A cells resulted in increased levels of MST2 (Fig. 2F). The level of p21, which is not a βTrCP substrate, CUL1 and SKP1 did not change upon βTrCP knockdown.

### Identification of a βTrCP binding site in MST2

The F-box protein subunits of SCFs, such as βTrCP, recognize their substrates via degradation motifs termed degrons (*40*). Most known SCF^βTrCP^ substrates contain the consensus sequence DpSGXX(X)pS (Fig. 3A), a biphosphorylated motif, that mediates the interaction of the substrate with the WD40 repeats of βTrCP. Although, MST2 does not contain a canonical βTrCP degron, we searched for sequences chemically similar to the highly conserved three amino-acid residues of the canonical degron (DpSG). Four putative tripeptide degrons are present in the human MST2 protein sequence: EDS (16-18), ESG (48-50), ESD (359-361), and EDG (375-377) (Fig. 3B). Noteworthy, similar sequences are found in other noncanonical degrons of experimentally validated SCF^βTrCP^ targets (*42–46*). We generated four MST2 constructs each containing a deletion for one of the four tripeptides and transfected them into HEK293T cells for co-IP experiments using an antibody against βTrCP. All 4 deletions affected MST2 binding to βTrCP, however deletion of the tripeptide EDG (375-377) almost abolished the interaction (Fig. 3C).

**Figure 3:**
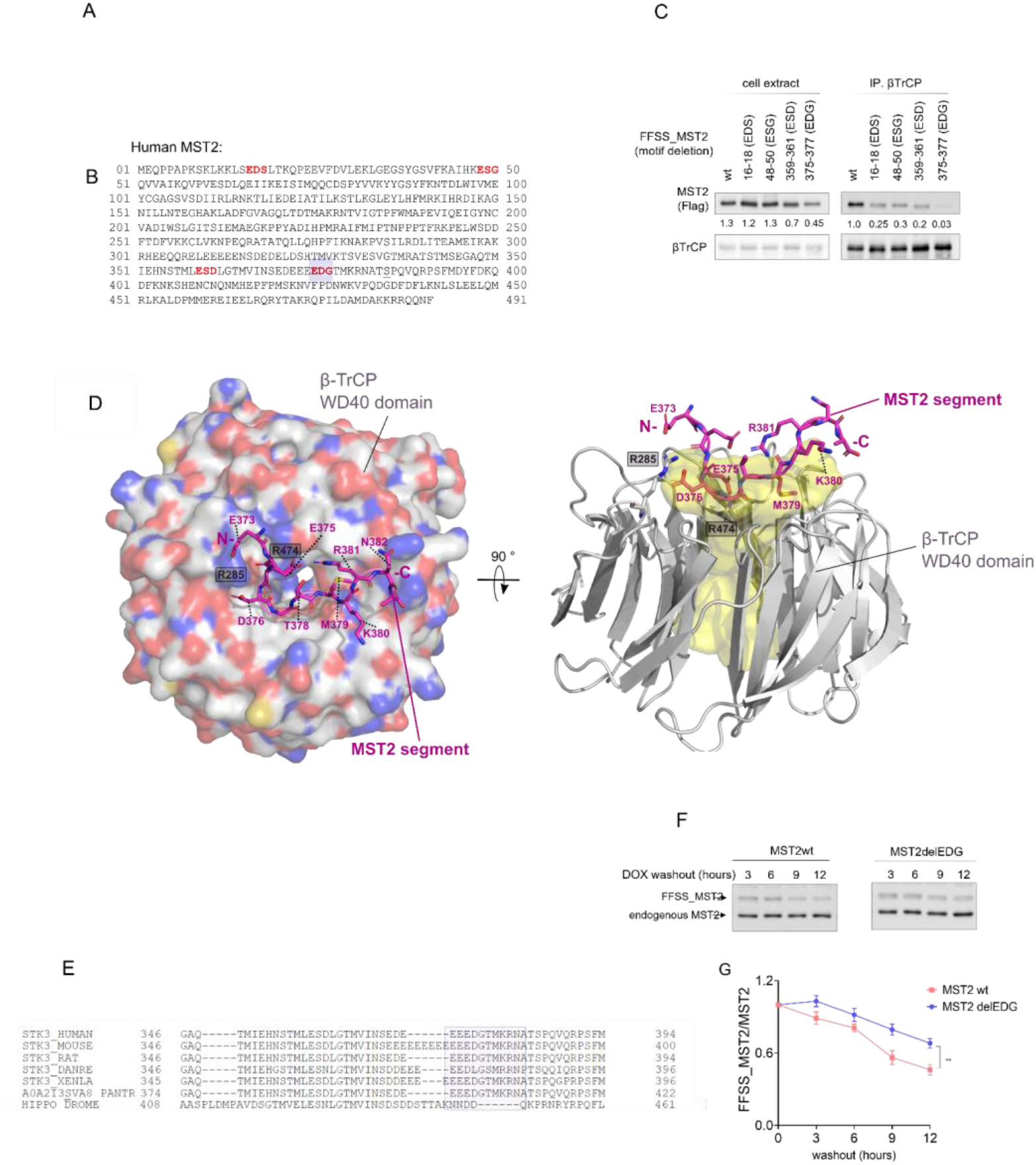
βTrCP binds to MST2 via a noncanonical degron. A) βTrCP consensus degron motif. The motif logo was generated with using MEME (http://meme-suite.org/tools/meme) according to Low et al (*5*). B) MST2 protein sequence. Four tripeptides chemically similar to DSG, the most conserved amino acid residues of the βTrCP canonical degron (Fig. 3A), are found (highlighted) in MST2 sequence: EDS (16-18), ESG (48-50), ESD (359-361) and EDG (375-377). C) Deletion of EDG (375-377) strongly impacts MST2 interaction with βTrCP. HEK293T cells were transfected with FFSS-tagged MST2 constructs which the tripeptides identified in (B) were deleted or a wild type (wt) construct. Cells were treated with MG132 (5 μM) for 1.5 hour and lysed. Endogenous βTrCP was immunoprecipitated, using a specific antibody and protein A/G-coated beads. Immunobloting for Flag was performed to detect FFSS-MST2. βTrCP was detected with the same antibody used for the immunoprecipitation. Total cell extracts (left) were also probed to assess the levels of the different constructs. Fold changes of FFSS_MST2 are shown under their respective immunoblots. D) Structural model of the MST2 EEEDGTMKRNA peptide segment bound to βTrCP WD40 domain obtained from 200-ns MD simulation. In the left, βTrCP WD40 domain surface is colored by element. The MST2 peptide is represented in magenta. MST2 residues are indicated by dashed lines and β-TrCP residues are indicated in boxes. In the right, the image shows a 90° rotated view of βTrCP WD40 domain represented as cartoon and the MST2 peptide by magenta sticks. The cavity occupied by MST2 peptide is rendered as yellow surface and was determined using the parKVFinder software (https://doi.org/10.1016/j.softx.2020.100606). Cavity detection was performed with the ligand adjustment mode using the following parameters: probe in of 1.4 Å, probe out of 15 Å, removal distance of 1.5 Å and ligand cutoff of 20 Å. MST2 peptide N- and C-terminals are indicated by letters. E) Interspecies alignment of MST2 protein sequence showing that the degron sequence EDGTMKRNA is conserved from Zebra Fish to Humans. DANRE = *Danio rerio* (zebrafish); XENLA = *Xenopus laevis* (African clawed frog); PANTR = *Pan troglodytes* (chimpanzee); and DROME = *Drosophila melanogaster* (fruit fly). F-G) MST2 with deleted EDG (375-377), MST2-delEDG, is more resistant to degradation than the wild type MST2 (MST2wt). F) HEK293T cells were transfected with FFSS-tagged MST2wt or MST2-delEDG constructs and treated for 24 h with doxycycline (Dox). After Dox washout, the cells were collected for protein extraction every 3 hours for 12 hours and submitted to immunoblotting with an antibody for MST2. Both ectopic and endogenous MST2 were detected as indicated in the figure. G) Quantification of MST2wt and MST2-delEDG after Dox washout. The intensity of FFSS-tagged MST2wt and MST2-delEDG bands was normalized to the intensity of the endogenous MST2. Data are show as mean ± SEM (N = 3). All cell culture assays in this figure were performed on regular cell culture polystyrene plates and flasks.

Because deletion of the MST2 EDG tripeptide produced the most prominent impairment in βTrCP interaction, we decided to model the MST2 segment that includes the EDG tripeptide in a complex with the βTrCP WD40 domain to investigate the structural basis that guides their recognition and interaction. The EDG tripeptide and most of its flanking residues are localized, as most degron motifs of E3-ligase targets, in a disordered region of MST2 (Supplementary Figure 2A). In comparison to the canonical βTrCP degron motif (Figure 3A), the first phosphorylated serine is substituted by an aspartic acid residue (D376) in MST2 EDG tripeptide, maintaining its negative charge feature (Figure 3A). Additional glutamic acid residues N-terminal to EDG could also help provide electrostatic forces that compensate the lack of phosphorylation. The second phosphorylated serine from the canonical degron (position +4 to +5 in relation to the first phosphorylated serine) is not present in the linear sequence of MST2, and two positively charged residues occupy this position instead.

Molecular dynamics (MD) simulation of an MST2 11-mer peptide (EE**EDG**TMKRNA), starting from the position of β-catenin 11-mer peptide crystallographic structure in complex with βTrCP (see methods), shows a small deviation at the EDG tripeptide region relative to its initial position (Supplementary Figure 2B-D). The stability in this region is a consequence of putative hydrogen bonds between D376 in MST2 (structural position similar to pS33 from β-catenin peptide) and R285 in βTrCP as well as E375 in MST2 (structural position similar to D32 from β-catenin peptide) and R474 in β-TrCP (Figure 3D and Supplementary Figure 2E). In MST2, E373 also contacts R285 through hydrogen bonds. Another similarity between MST2 and β-catenin peptide binding is the presence of a hydrophobic residue (M379 of MST2 and I35 in the β-catenin peptide) pointing towards the βTrCP β-propeller channel (Figure 3D and Supplementary Figure 2E).

During the simulation, K380 in MST2 is displaced from its initial position to avoid repulsive contact with R431 in βTrCP (Supplementary Figure 2B), which anchors the second phosphorylated serine in canonical degrons (Supplementary Figure 2E). The presence of βTrCP positive charges favors K380 in MST2 to be exposed to solvent. This is an interesting finding, since an *in silico* analysis predicted that K380 could be ubiquitinated (Supplementary Figure 3A).

Protein degradation motifs are commonly conserved across species (*42*). We performed interspecies alignment of MST2 and found that the MST2 EE**EDG**TMKRNA peptide is conserved from zebra fish to Mammalians, but it is not present in the ancestor orthologue Hippo in Drosophila (Fig. 3E).

We then assessed whether deleting the tripeptide EDG would alter MST2 susceptibility to degradation. For this, we cloned MST2-wild type (wt) or MST2-delEDG in a tetracycline inducible expression vector transfected them into HEK293T cells. After Doxycycline (Dox) treatment, both vectors showed robust protein expression (Supplementary Figure 4). A twelve-hour pulse chase-like experiment - induction with Dox followed by its removal - confirmed that MST2-delEDG was indeed more resistant to degradation than MST2-wt (Fig. 3F-G). Altogether, our experimental and computational analyses revealed that MST2 can bind to βTrCP WD40 domain through the noncanonical EDG site, but maintaining structural and chemical characteristics of canonical degrons, such as the β-catenin degron.

**Figure 4:**
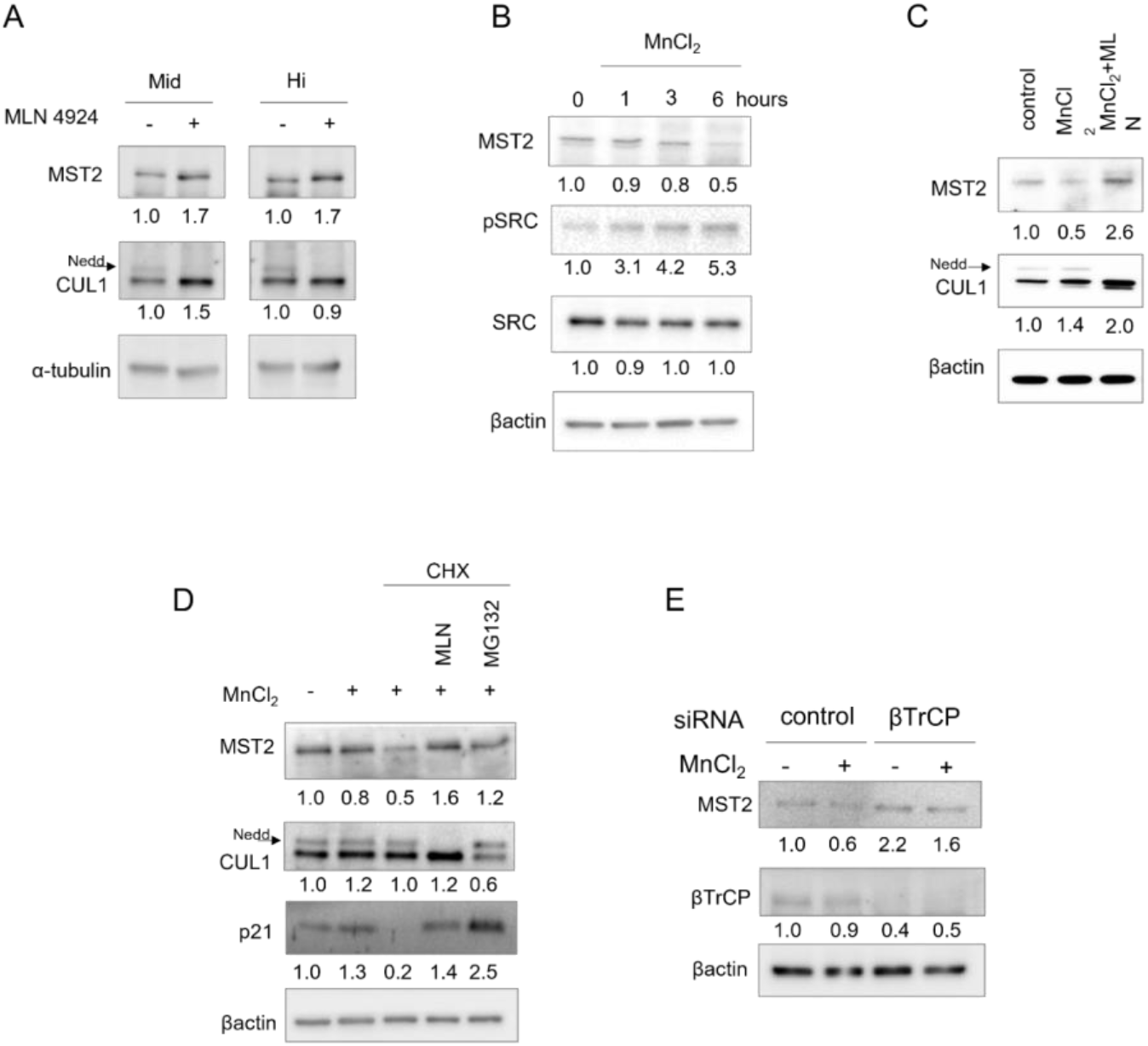
ECM stiffness and integrin activation triggers MST2 degradation. A) Immunoblots of MCF10A cells cultured on Low and Hi rigidities and treated with CUL1 neddylation inhibitor (MLN4924). After MLN4924 (MLN) treatment, MST2 increases in both degrees of rigidity. Note that the band referring to CUL1 neddylation (indicated by an arrow) disappears in the presence of MLN4924 indicating that the inhibition was efficient. α-tubulin was used as loading control. B) Immunoblots of MCF10A cells treated with MnCl_2_ (1 mM) for 1, 3 and 6 hours. SRC phosphorylation was detected to assess integrin signaling activation. β-actin was used as loading control. C) Immunoblots of MCF10A cells pretreated with MLN for 1.5 hours and then treated or not with MnCl_2_ (1 mM) for 6 hours. β-actin was used as loading control. Note that the band corresponding to neddylated CUL1 (indicated by an arrow) disappears in the presence of MLN4924 indicating efficient inhibition. D) Immunoblots of MCF10A cells treated with MnCl_2_ for 3 hours, with the protein synthesis inhibitor cycloheximide (CHX) for 4 hours, with MLN4294 and MG132 for 1.5 hours. The different combinations of treatments are indicated in the figure panel. Note that the band referring to CUL1 neddylation (indicated by an arrow) disappears in the presence of MLN4924 indicating that the inhibition was efficient. MST2 levels robustly decrease in the presence of MnCl_2_ and CHX, but inhibition of the proteasome or neddylation reverses this effect. Immunoblotting for p21 was used as a positive control to assess efficiency of CHX, MLN4294 and MG132 treatments. β-actin was used as loading control. E) Silencing of βTrCP in MCF10A cells results in MST2 accumulation, despite integrin signaling hyperactivation. Cells were cultured in the presence of siRNA for βTrCP or a non-targeting siRNA (siControl) and treated or not with MnCl_2_ for 6 hours. Immunoblots confirming that silencing of βTrCP was efficient and resulted in increased MST2 levels even in the presence of MnCl_2_. β-actin was used as loading control. (A-E) Fold changes of protein levels and phosphorylation of SRC (pSRC/total SRC) are shown under their respective immunoblots.

### Stiff ECM substrates and activation of integrin signaling triggers MST2 degradation

We sought at investigating the signaling pathways activating MST2 degradation in stiffer substrates. First, we confirmed that inhibition of CUL1 neddylation resulted in accumulation of MST2 in MCF10A cells cultured on stiff ECM (Mid and Hi hydrogels) (Fig. 4A). Because stiff microenvironments trigger integrin signaling (*4, 9, 47, 48*), we asked whether modulation of integrin signaling would result in changes of MST2 levels. We mimicked integrin hyperactivation by treating MCF10A cells grown on polystyrene plates with MnCl_2_. Integrins drastically increase affinity to their ligands when bound to Mn^2+^ (*49, 50*). A time-course experiment showed that MST2 decreases in response to Mn^2+^ (Fig. B).

Inhibition of CUL1 neddylation prevented the effects of integrin hyperactivation with Mn^2+^ (Fig. 4D). A 3-hour treatment with Mn^2+^ (Fig. 4E) resulted in 20% reduction of MST2 levels, but a co-treatment with cycloheximide (CHX), a protein synthesis inhibitor, resulted in robust MST2 decrease, similar to the response shown by p21, a well-established ubiquitin-proteasome system target. Inhibition of CUL1 neddylation or proteasome activity reverted the effects on MST2 content caused by the co-treatment (Fig. 4E). To assess the direct association between integrin activation and MST2 ubiquitination by SCF^βTrCP^, we silenced βTrCP in combination with integrin hyperactivation by Mn^+2^. In fact, forcing decreased levels of βTrCP abrogated the effects of integrin signaling on MST2 (Fig. 4F).

FAs are large protein complexes composed of kinases, FAK and/or ILK, integrins and scaffold proteins that mechanically and biochemically link the ECM to non-cytoskeleton and cytoskeleton proteins resulting in a multitude of cellular outputs (*52, 53*). Our hypothesis was that pharmacological perturbation of the main FA kinases (FAK and ILK) in MCF10A cells would impact MST2 proteostasis. Immunoblots for phospho-AKT and phospho-FAK confirmed that ILK and FAK were inhibited. Surprisingly, kinetical studies showed that only ILK pharmacological inhibition resulted in higher amounts of MST2, whereas FAK inhibition did not alter the level of MST2 (Fig. 5A-B). To further verify the involvement of ILK in MST2 proteostasis, we inhibited ILK in conjunction with integrin hyperactivation by Mn^2+^. ILK inhibition resulted in increased levels of MST2 (Fig. 5C), and decreased MST2-βTrCP binding in MCF10A cells, as assessed by Co-IP of endogenous MST2 (Fig. 5D). Importantly, this latter experiment reinforces the robustness of our findings since it shows the binding of endogenous MST2 and βTrCP in relevant cell model.

**Figure 5:**
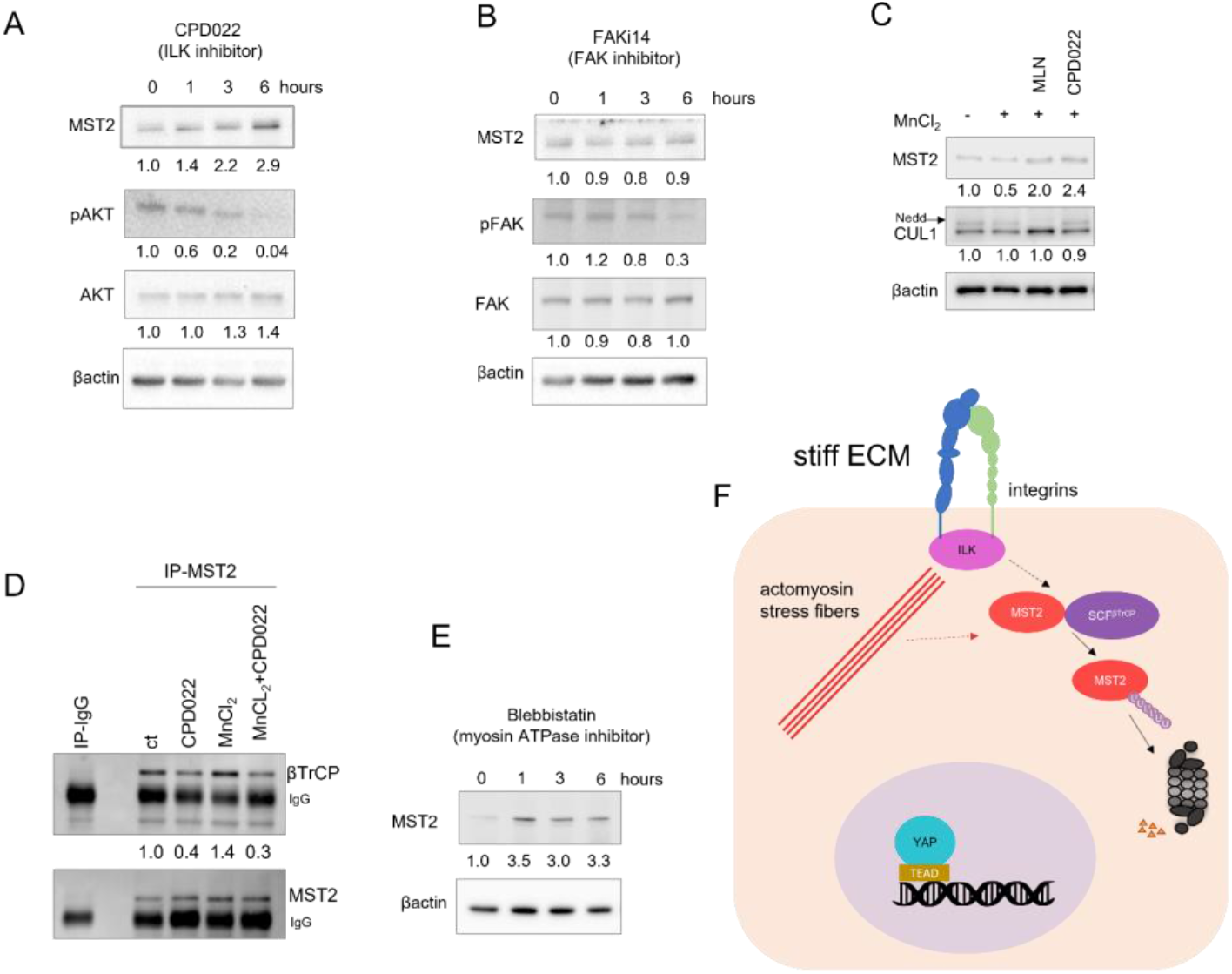
Perturbation of ILK and the actomyosin cytoskeleton regulates MST2 levels: A) Immunoblots of MCF10A cells treated with an Integrin-Linked Kinase (ILK) inhibitor (CPD022) for 1, 3 and 6 hours. AKT phosphorylation was detected to assess the efficiency of ILK inhibition. β-actin was used as loading control. B) Immunoblot for MST2 of MCF10A cells treated with FAK inhibitor (FAKi 14) for 1, 3 and 6 hours. FAK phosphorylation was detected to monitor its inhibition. β-actin was used as loading control. C) Immunoblots of MCF10A cells treated with CPD022 or MLN4294 in combination with MnCl_2_ for 6 hours. Both inhibition of CUL1 neddylation and ILK reverses the effect of integrin hyperactivation. β-actin was used as loading controls. Note that the band corresponding to CUL1 neddylation (indicated by an arrow) disappears in the presence of MLN4924 indicating that the inhibition was efficient. D) Co-IP revealed that the interaction of endogenous MST2 and βTrCP decreases after ILK inhibition, despite integrin hyperactivation. MCF10A cells treated with DMSO (ct), CPD022, MnCl_2_ or CPD022 concomitantly with MnCl_2_ for 6 hours. Co-IP was performed with an MST2 antibody or a control IgG, followed by incubation with protein A/G-coated beads. βTrCP and MST2 were only detected in the MST2 immunoprecipitants. MST2 was detected to confirm the Co-IP. Arrows indicate bands corresponding to IgG that were detected by the secondary antibody. E) Immunoblot for MST2 of MCF10A cells treated with a myosin ATPase activity inhibitor (Blebbistatin, 2.5 μM) for 1, 3 and 6 hours. β-actin was used as loading control. (A-E) Fold changes of protein levels and protein phosphorylation (phosphorylated/total) are shown under their respective immunoblots. F) Schematic depicting the pathway of MST2 degradation induced by ECM stiffness. In a stiff ECM, hyperactive integrin signaling results in ILK activation and actomyosin contraction leading to ubiquitination of MST2 by SCF^βTrCP^ and consequent MST2 degradation via proteasome 26S. This mechanism influences the level of YAP in the nucleus. All cell culture assays presented in this figure were performed in polystyrene cell culture plates.

Stiff substrates elevate the activity of growth factor activated pathways in cancer, fibrosis and other diseases (*54, 55*). For example, PIK3-AKT is a major growth promoting pathway that can be amplified by integrin activation. However, pharmacological inhibition of PI3K or AKT did not affect MST2 concentration (Suppl. Fig 5A-B). Because stiff microenvironments enhance actomyosin contraction, we pharmacologically inhibited myosin-II with blebbistatin, an inhibitor of myosin ATPase activity, and measured the level of MST2. We could observe a relatively transient increase in MST2 levels after myosin contraction inhibition by both drugs (Fig. 5E-F), indicating that intracellular tension stored in the actomyosin cytoskeleton plays a role in MST2 proteostasis. This set of experiments revealed that ILK directly links integrin activation to MST2 degradation control in an actomyosin-cytoskeleton manner (Fig. 5G).

## Discussion

In addition to playing crucial roles in tissue development and regeneration, Hippo kinases can be considered tumor suppressors (*56*). Here, we uncovered the molecular mechanism by which MST2, one of the core Hippo kinases, degradation is regulated (Fig. 5G). We not only identified that SCF^βTrCP^ ubiquitinates MST2 for proteasomal proteolysis and mapped a βTrCP binding motif (degron) in MST2, but also described a pathway based on integrin and ILK activation that triggers MST2 degradation in breast epithelial cells.

The cell microenvironment provides active and passive mechanical cues that elicit biochemical signaling through the cytoskeleton and kinase cascades regulating a milieu of cellular processes such as cell proliferation (*57*), invasion (*4*), metabolism (*55*) and differentiation (*58*). A crucial signal is the ECM rigidity. This is initially sensed by integrins which are intracellularly linked to the actomyosin cytoskeleton, which in response to stiffness, increase cell contractility. Tension built in contractile actomyosin reciprocally activates the assembly of FA (*26, 57*). YAP/TAZ accumulate in the nuclei of epithelial cells, but compelling evidence has shown that YAP/TAZ exert at least part of their mechanoregulatory activity by directly inducing the expression of genes encoding for FA proteins (*59*). Thus, reinforcing a feed-forward activation cycle where a stiff ECM increases YAP/TAZ nuclear activity and YAP/TAZ transcriptional control activity results in increased FA assembly. We indeed observed that *STK3* knock-down affects proliferation and cell shape regardless of rigidity. YAP ratios at high rigidity are not affected, but this is likely because they are already saturated at their maximum values. Furthermore, the rigidity dependence of YAP after *STK3* silencing remains very robust. Potentially, the effects we observed might be downstream of YAP: YAP activity feedbacks to regulate focal adhesions and stress fibers, so these responses could be generally primed by preventing phosphorylation by MST2. This hypothesis will be addressed in future studies from our laboratory.

There are conflicting data in the literature as to whether the mechanical cues that influence YAP/TAZ localization and activity are mediated by Hippo kinases (*10, 60, 61*). This might be attributed to the use of different cell lines, type of ECM molecule used as substrate, hydrogel preparation and degrees of stiffness. Nevertheless, in the present work the outputs in our MST2 silencing assays were different depending on the cellular process assessed: MST2 knock down resulted in increased proliferation in all three degrees of rigidity while cell shape and F-actin arrangement were only perturbed in stiffer substrates indicating that regulation of cell proliferation and cytoskeleton/cell shape might be independently regulated in physiological stiffness and also that YAP/TAZ regulation of FA assembly might occur only in stiffer ECMs. Although our data clearly shows that MST2 level is mechanoresponsive, other complementary factors could act on YAP/TAZ as previously reported (*10, 62*). Also, MST2 can phosphorylate a vast number of substrates (*63*) and play roles nonrelated to its kinase activity ((*64*). Therefore, part of the phenotypic changes due to MST2 modulation might not relate to regulation of YAP/TAZ but by action of other effectors. In this context, there is evidence showing that cytoskeletal disruption activated MST as well as JNK1 in an MST-dependent manner (*65*). Inhibition of actin cytoskeleton with tat-C3 resulted in higher MST activity (*65*).

A series of biochemical assays, using ectopic protein expression in HEK293T cells, unveiled that SCF^βTrCP^ tags MST2 for degradation after binding to a non-canonical degron in MST2. Crucially, by using a HMEC and probing for endogenous proteins, we not only confirmed the robustness of our findings - corroborating that the results from assays with HEK293T cells were not specific to this cell line neither an artifact resulting from nonphysiological overexpression of protein constructs - but also showed that integrin hyperactivation increases SCF^βTrCP^ and MST2 binding. (Fig. 2G). We also observed that inhibition of ILK and dissipation of tension built in actomyosin stress fibers caused MST2 accumulation in HMEC and inhibited SCF^βTrCP^ and MST2 binding, indicating that MST2 degradation is directly dependent upon the integrin-ILK axis activation.

The βTrCP degron in MST2 does not contain serine amino-acid residues as the canonical degron, raising the question of how the MST2 and βTrCP interaction is molecularly regulated. Nevertheless, the chemical properties of the MST2 tripeptide EDG are similar to the conserved DpSG in the canonical degrons and the *in silico* studies showed that an 11-mer MST2 peptide that includes the EDG tripeptide stably binds to the βTrCP, despite the fact that the segment does not present a C-terminus negative amino acid that would substitute for the second phosphorylated serine in the degron. It remains, thus, to be elucidated whether phosphorylation in other regions of MST2 or other posttranslational modifications would influence MST2 and βTrCP binding. Moreover, it will be important to pinpoint which molecule is linking integrin-ILK activation to MST2 and βTrCP. There are, indeed, other instances in the literature of degrons that do not follow the DpSGXX(X)pS rule. For example, one of the WEE1 degrons (SWEEEGFGSS) (*43*) and the CDH1 degron (SSPDDGNDVS) (*44*) do not bear the first serine found in the canonical degron. And, there is at least one report of a degron (the CDC25B degron; TEEDDGFVDI) analogous to the MST2 degron we identified, that does not contain any serine (*45*).

Frequently, tumors are stiffer than normal tissues eliciting integrin hyperactivation (*4, 8, 48*). We propose that engagement of integrin-ILK in focal adhesion arrays in stiffer ECM, could result in decreased levels of MST2 due to enhanced protein degradation and this process could cooperate with other aberrant signals in reinforcing malignant phenotypes in tumor microenvironments. We also suggest that the pathway, integrin-ILK-MST2, might consist in a novel molecular target for treating breast tumors, and potentially other types of cancers, that display increased density mammograms and low levels of Hippo activity. Based on the degron we identified, a possible strategy would be developing peptides or small molecules that would block MST2 degradation by blunting the anchoring of βTrCP to MST2. MST2 bears a noncanonical βTrCP degron, hence precisely targeting the βTrCP-MST2 interaction interface could reduce the chance of undesirably off-targeting SCF^βTrCP^ substrates with opposing biological functions. For instance, although YAP and TAZ are substrates of SCF^βTrCP^ their degrons differ considerably from the one we described for MST2. Finally, we believe that our work unveiled a detailed mechanism regulating MST2 levels and this is expected to have wide ramifications for cell tensional homeostasis and offer a novel molecular opportunity for therapeutic strategies.

## Materials and Methods

### Cell lines and cell culture media

Non tumoral breast epithelial cell lines MCF10A (Gently donated by Mina J. Bissell -Lawrence Berkeley National Laboratory) and MCF12A (Gently donated by Carlos Frederico Martins Menck, from Instituto de Ciências Biomédicas-USP) were cultured in Dulbecco Modified Eagle’s/ HAM F-12 medium (1:1, DMEM/F-12, Gibco, #12-500-062), supplemented with 5% horse serum (Gibco # 16050-122), Insulin (10 μg/mL, Sigma-Aldrich, #I5500), Hydrocortisone (10 μg/mL, Sigma-Aldrich # H4001), Choleric toxin (1 μg/mL, Sigma-Aldrich, #C8052) and Epidermal Growth Factor (20 ng/mL, Sigma-Aldrich, #E4127).

Breast cancer epithelial cell lines MCF7 and MDA-MB231 (both gently donated by Mina J. Bissell - Lawrence Berkeley National Laboratory), and MDA-MB468 (purchased from American Type Culture Collection, ATCC, #HTB-132) cell lines were cultured in DMEM high glucose (Gibco, #12100046) supplemented with 10% of fetal bovine serum (FBS, Gibco #12657-029); BT474 and SKBR3 (both purchased from ATCC #HTB-20 and #HTB-30, respectively) were cultured in RPMI (Gibco #11875093) supplemented with 10% FBS, Hyclone non-essential amino acids (0.1 mM, Thermo #SH3023801), L-glutamine (2 mM, Gibco, #25030081) and sodium pyruvate (1 mM, Gibco, #11360070).

Human Embryonic Kidney (HEK) 293T cells (purchased from ATCC, #CRL-11268) were cultured in Dulbecco’s Modified Eagle’s Medium (DMEM High Glucose, Thermo Fisher) supplemented with 10% Fetal Bovine Serum (Gibco), Hyclone non-essential amino acids (0.1 mM, Thermo), L-Glutamine (2 mM, Gibco) and sodium pyruvate (1 mM, Gibco).

### Cell culture on polyacrylamide hydrogels

The polyacrylamide hydrogels with tunable mechanical properties were made in three elastic modulus levels: 0.48 kPa (3% acrylamide, 0.06% bis-acrylamide), 4.47 kPa (5% acrylamide, 0.15% bis-acrylamide) and 40.40 kPa (8% acrylamide, 0.48% bis acrylamide), as previously described (*29*). The hydrogels were coated with fibronectin-1 (10 μg/ml) by using the crosslinker Sulfo-SANPAH (Pierce Biotechnology, # A35395). MCF10A cells were seeded at a density of 3 × 10^4^ cells per cm^2^ in complete medium and kept in culture for 72 hours.

### cDNA constructs, plasmids and transfection

All plasmids for protein expression used in this study are listed in Table S1. *STK3* (accession number NM_006281.4) ORF was amplified from MCF10A human mammary cells using specific primers (listed in Table S2) with Q5 High-Fidelity 2X Master Mix (New England BioLabs, # M0492), following the manufacturer’s instructions. The MST2 ORF was inserted in the pCDNA3.1_Flag2x _Strep2x vector using StickyEnd ligase (New England Biolabs, # M0370L). The plasmids pCDNA3.1_Flag_ β-TrCP, pCDNA3.1_Flag_ βΔFbox and pCDNA3.1_myc_ubiquitin were described previously (*66, 67*). To obtain the MST2 constructs with the tripeptide deletions, we used the pCDNA3.1_Flag2x _Strep2x _MST2 plasmid as a template, specific primers (MST2_BAMHI_Fw and MST2_XHOI_Rv, listed in Table S2) and the site directed deletions and amplification were performed using Pfu turbo DNA polymerase (Agilent, #600250) following manufacturers protocol. All constructs’ sequences were confirmed by Sanger sequencing (GeneWitz, NJ) and were transfected in HEK293T cells using the Lipofectamine 3000 reagent (Thermo Fischer # L3000008).

To generate inducible vectors for MST2 wild type and del EDG we amplified the Flag2x _Strep2x fused to the MST2 constructs (wild type and del_EDG) with Q5 High-Fidelity Polymerase (New England BioLabs, # M0492) using specific primers (EcoRI_Flag_Fw and MST2_AgeI_Rv, listed in table S2) and inserted the 2xFlag_2xStrep_MST2 constructs (wild type or del_EDG) in the inducible vector pLVX (Clontech, #631847).

### Gene silencing by siRNA

The following pre-designed and validated duplexes siRNA oligos (Ambion # 4392420): *STK3* (s13567), *βTrCP* (s17109), and non-targeting (siRNA #1, AM4635) were used. Oligo duplexes were transfected using Lipofectamine RNAiMax (ThermoFisher Scientific # 13778030) according to the manufacturer’s instructions, using 200 nM of each oligo. The durations of transfection are indicated in the figures and/or figure legends.

### Treatment with inhibitors and blocking peptides

Inhibitions of Cullin (CUL1) neddylation or proteasome activity were achieved by treating cells with 2.5 μM MLN4924 (Cayman Biochemicals # 15217) or 5 μM MG132 (Peptides International # IZL-3175-v) for the durations indicated in the figure legends. To assess the protein stability of MST2, we inhibited global protein synthesis with cycloheximide (CHX, 100 μg/mL, Sigma Aldrich # C4859).

Integrin activation was induced in MCF10A cells with 1 mM MnCl_2_ (Sigma Aldrich # M1787) treatment for the durations indicated in the figure legends (*68*). To screen the pathways mediating MST2 degradation, MCF10A cells were treated for 1, 3 and 6 hours with the inhibitors for ILK (1 μM CPD022, Calbiochem, #407331), FAK (5 μM FAK inhibitor 14, Tocris # 3414), PI3K (30 μM LY294002, Gibco # PHZ1144), AKT (20 μM MK2206, Cayman Biochemicals #1032350-13-2) and myosin ATPase activity inhibitor (5 μM Blebbistatin, Tocris #1852).

### MST2 stability assay using an inducible protein expression system

HEK293T cells were transiently transfected using Lipofectamine 3000 (ThermoFisher Scientific) based on the manufacturer’s recommendation with wild type MST2 or MST2_delEDG subcloned in pLVX_TetOne-Puro (Clontech # 631849). Cells were cultured in presence of Doxycycline (Dox, 500 ng/mL Sigma Aldrich, #D9891) to induce MST2 and MST2_delEDG expression. After 24 hours, Dox was washed with PBS 3 times for 30 minutes and complete medium was added. Cells were collected for protein extraction 3, 6, 9 and 12 hours after Dox removal.

### Protein extraction, immunoprecipitation, SDS-PAGE and Immunoblot

For total cell lysate protein extraction, samples were lysed using modified RIPA buffer (Pierce) supplemented with protease (Sigma-Aldrich) and phosphatase inhibitors (Sigma-Aldrich). Protein concentration was determined using the DC Protein Assay kit (BioRad, #5000111). Protein lysate was resuspended at a 0.5 μg/μL concentration in Laemmli sample buffer (0.1% 2-Mercaptoethanol, 0.005% Bromophenol blue, 10% Glycerol, 2% SDS, 63 mM Tris-HCl), heated at 95°C for 5 minutes and kept at −20 °C.

For the Co-immunoprecipitation (Co-IP) cells were lysed in immunoprecitation lysis buffer (25 mM Tris pH 8.0, 150 mM NaCl, 10% glycerol, 1 mM EDTA, 1 mM EGTA, 1 mM 1,4-Dithiothreitol (DTT) and 0.1% NP-40) containing protease (Sigma-Aldrich) and phosphatase inhibitors (Sigma-Aldrich). The insoluble fraction was removed by centrifugation (20,000 x g x 15 min at 4°C). Protein concentration was determined using the DC Protein Assay kit (BioRad #5000111). For both MST2 and βTrCP immunoprecipitation, we incubated 2 mg of protein extracts with specific antibodies (listed in Table S3) overnight at 4°C. Subsequently, we added Protein A/G magnetic beads (Pierce, # 88802), incubated at 4°C for 2 hours, and proceeded with extensive washing using the lysis buffer. After the final wash, the beads with the immunoprecipitant were resuspended with 1x Laemmli sample buffer, heated at 95°C for 5 minutes and kept at −20 °C.

The flag-tag immunoprecipitations were carried out using FLAG-M2 coated agarose beads (Sigma-Aldrich #F2426) for 2 hours at 4 °C. The beads were then extensively washed in lysis buffer and elution was carried out with 50 μL of 3×FLAG peptide (Sigma-Aldrich #F4799), the supernatant was separated from the beads by centrifugation, resuspended in the same volume of 2x Laemmli sample buffer, heated at 95°C for 5 minutes and kept at −20 °C.

Total protein extracts or immunoprecipitants were resolved in SDS-PAGE followed by immunoblot. In brief, 10 μL (5 μg) of protein sample and 20 μL of IP samples were heated at 95°C for 5 minutes and loaded into 10% tris-glycine polyacrylamide gels. Resolved proteins were transferred to PVDF membrane (0.45 μm, Millipore, #05317) followed by 1-hour incubation in a blocking buffer (TBS, 0.5% Tween-20 with 1% BSA). Membranes were incubated in buffer (TBS, 0.5% Tween-20) containing primary antibodies (listed in Table S3) overnight., and finally incubated with horseradish peroxidase (HRP) for 1 hour at room temperature in the blocking buffer. HRP was detected by Pierce SuperSignal detection kit (Thermo-Fisher Scientific, #A45916) and the chemiluminescence signal was captured with a ChemiDoc MP Imaging System (BioRad). All immunoblots for probing total cell lysates were performed twice.

### RNA extraction, reverse transcription, and quantitative polymerase chain reaction (RT-qPCR)

Total RNA was extracted using RNeasy kit (Qiagen #74106) following the manufacturer’s guidelines. The RNA 260/280 and 260/230 ratio and the concentration were measured with a Nanodrop (Thermo) UV-spectrophotometer. The cDNA was generated using 1 μg of total RNA using the kit First-Strand cDNA Synthesis (Thermo Scientific # 18080051) using oligoDT and random hexamer primers, following the manufacturer’s instructions. For the qPCR, 10-μL reactions were performed in triplicates with 25 ng of cDNA using 2x SYBR Green PCR Master Mix (Applied Biosystems # 4364346) and specific primers (listed in table 2) in MicroAmp Optical 96-Well Reaction plates (Applied Biossystems #N8010560). The reactions were performed in the AB-9500 system (Applied Biosystems) real-time thermocycler. GAPDH was used as an endogenous control and mRNA fold-changes were calculated according to the Pfaffl method (*69*).

### 5-ethynyl-2’-deoxyuridine (EdU) incorporation assay

The Click-iT® EdU kit (Life Technologies, # C10639) was used for measuring the percentage of cells in S-phase in different conditions cultured on the hydrogels. Only DNA-replicating cells incorporate the base analog EdU. Cells were incubated with 20 μM EdU for 1 hour, and fixed with 4% PFA (Electron Microscopy Sciences, #15714) in PBS. EdU incorporation was detected following the manufacturer’s instructions as previously described (*70*). The cells were then counterstained with 4’6-diamidino-2-phenylindole (DAPI, 0.5ug/mL, Sigma-Aldrich #D9542) for 10 minutes, the slides/ coverslips were mounted with Prolong® Diamond (Thermo Scientific # P36965) and analyzed with a Leica DMi8 wide-field microscope (Leica Microsystems) equipped with a 40x objective.

### Immunofluorescence and phalloidin staining

The cells were fixed with 4% paraformaldehyde (PFA) (Electron Microscopy Sciences) in PBS. Residual PFA reactivity was quenched with PBS–25 mM glycine and cells were permeabilized with 0.5% Triton X-100 in PBS for 30 minutes. The cells were then incubated in blocking buffer 1% Bovine Serum Albumin (MP Biomedicals, #160069) and 5% goat-serum (Gibco #6210-064) in PBS for 1 hour at room temperature, followed by overnight incubation with primary antibody (listed in table S3) in blocking buffer. Cells were then incubated in blocking buffer with Alexa 488 or Alexa 555 conjugated secondary antibodies (listed in table S3) for 45 minutes, counterstained with DAPI for 10 minutes and mounted with Prolong® Diamond (Thermo Scientific #P36970) and analyzed with a Leica DMi8 wide field microscope. Image stacks were captured with a 63X/1.4 NA oil objective (Leica) for YAP detection. We used the optimized step intervals along the Z-axis to capture 3D images and the deconvolution of the acquired images was performed in the Leica Application Suite-X software using the tool blind deblur with 10 iterations (Leica Microsystems).

Superresolution fluorescence microscopy images of cells stained with phalloidin-Alexa 488 (1:50, Thermo Fisher Scientific #A12379) and vinculin antibody (1:500, Sigma #V9131) were obtained with an LSM-880 confocal microscope equipped with the AiryScan system (Carl Zeiss) using a 63x/1.4 NA oil objective lens and optimized acquisition settings. All imaging acquisition parameters were kept constant for each experiment.

### Image analysis

Fluorescence microscope and gel image quantification were performed using the ImageJ software (2.0.0-rc-68/1.52e; Java 1.8.0_66 64-bit). For the relative quantification of the protein levels of the immunoblot experiments, and each band integrated optical density was determined using the gel analyzer application in Image. The loading control (β-actin or α-tubulin, depending on the experiment) was also determined and the fold change was determined as the ratio between target protein level and loading control. For phosphorylated proteins, we used the ratio phosphorylated protein/ total protein/loading control.

For quantification of YAP staining, we used a customized ImageJ macro (available upon request). In brief, the DAPI channel was converted to a binary mask to delimit the nuclei and the analyze particles tool in ImageJ tool was used to generate selections (outlines) of individual nuclei. The imageJ tool *enlarge* was used to produce an outer and an inner ring by concentrically expanding or shrinking the nuclear. To determine the intensity of YAP level in the cytoplasm, the YAP intensity inside the nuclear outline was subtracted from the YAP staining in the outer ring. Finally, to obtain the YAP cytoplasm to nucleus ratio for each cell, the intensity of YAP signal inside the inner ring was divided by the intensity of YAP in the cytoplasm of the same cell.

To quantify the EdU assay, the number of total and Edu-stained cells was obtained by using the *cell counter* application in ImageJ and the percentages of EdU-positive cells were determined.

For quantification of F-actin directionality, we used the *directionality* plugin (https://imagej.net/Directionality) in ImageJ was used. For this, superresolution images of individual cells stained with phalloidin were aligned in respect to the cell’s major axis.

### *In vitro* ubiquitination assay

We co-transfected HEK293T cells with plasmids containing myc-tagged ubiquitin, 2x Flag 2x Strep MST2 and Flag βTrCP (listed in table1) for 24 hours. Three hours prior to collection, we added 5 μM MG132, a proteasome inhibitor. Protein extraction was performed under denaturing conditions (50 mM TRIS, 150 mM NaCl, 1 mM EDTA, 1% SDS) containing protease (Sigma-Aldrich #11836145001) and phosphatase inhibitors (Sigma-Aldrich #P5726-5ML) and the deubiquitinase inhibitor N-ethylmaleimide (1 μM, Sigma-Aldrich, #E3876). Samples were boiled at 98 °C for 10 minutes and sonicated 3 times for 10 seconds on ice and the insoluble fraction was removed by centrifugation. Next, 2xFlag 2xStrep MST2 was purified by affinity, using Strep-Tactin Superflow beads (IBA # 2-1206-025) in immunoprecipitation lysis buffer (50 mM TRIS, 150 mM NaCl, 2.5 mM EDTA, 1% Glycerol, 0,1% NP40) for 2 hours at 4°C, beads were extensively washed in lysis buffer and elution was carried out with Strep-Tactin Elution Buffer containing Desthiobiotin (IBA # 2-1000-025).

### Statistics

Statistical analyses were performed using GraphPad Prism 7.0. The number of samples analyzed and the number of times that the experiments were performed are indicated in the respective figure legends. All the data were submitted to a normality test. Nonparametric data were submitted to Kruskal-Wallis test, while parametric data were submitted to ANOVA followed by Dunnett test. Differences were considered significant when the p-value was less than 0.05. All graphs were generated with GraphPad and the data are presented as mean ± SEM.

### Molecular modeling and dynamic simulation and protein sequence alignment

Human MST2 secondary structure was predicted using the PsiPred server (http://bioinf.cs.ucl.ac.uk/psipred/) (*71*). An initial model of the MST2 segment that comprises EDG tripeptide (EEEDGTMKRNA) bound to β-TrCP WD40 domain was built using the main chain atoms coordinates of β-catenin peptide in a crystallographic complex with β-TrCP (PDB ID: 1P22) and changing the sidechains to match MST2 sequence. This process was performed using YASARA software (http://www.yasara.org/). Only βTrCP WD40 domain was used in modeling and simulations.

The molecular dynamic (MD) simulations were performed using YASARA with AMBER14 forcefield. For all simulations, the structures were solvated in a 10 Å cubic cell around all atoms, neutralized and submitted to energy minimization. The production runs were performed at 298 K and pH 7.4 with 2*2.5 fs timestep. All bonds and angles involving hydrogens were constrained using LINCS algorithm and an 8 Å cutoff was used for Van der Waals and Coulomb force calculation and long-range Coulomb interaction were calculated using the Particle Mesh Ewald algorithm. Simulation snapshots were saved each 250 ps. The production run of MST2 11-mer peptide EEEDGTMKRNA was performed for 200 ns and the last frame was submitted to energy minimization in YASARA and evaluated.

For MD simulations analysis, simulation snapshots were aligned using the βTrCP WD40 domain first snapshot as reference in VMD (*72*). RMSD (root-mean-square deviation) and RMSF (root-mean-square fluctuation) were calculated for the MST2 peptide using the R Bio3D package (*73*).

For alignment of MST2 sequence from different species, we used the multi sequence alignment tool from the software Jalview 2.11.1.3 (https://www.jalview.org/).

## Acknowledgements

This work was supported by Fundação de Amparo à Pesquisa do Estado de São Paulo (FAPESP) Young Investigator Award (14/10492-0), by a FAPESP grant (2019/26767-2) by the Conselho Nacional de Desenvolvimento Científico e Tecnológico (CNPq-Universal 444597/2014-0) and Coordenação de Aperfeiçoamento de Pessoal de Nível Superior (CAPES, Finance Code 001). MP is an investigator with the Howard Hughes Medical Institute and his lab is partially funded by grants from the National Institute of Health (R01-CA76584 and R35-GM136250). APZPF is a FAPESP Postdoctoral fellowship awardee (14/25832-1 and 17/18641-3). AMRS is funded by CAPES PhD-scholarship (PROEX), PFR is a FAPESP PhD-scholarship recipient (17/18067-5), MCSB is a FAPESP PhD-scholarship recipient (2017/25437-3) and ACM is a CNPq PhD-scholarship recipient (14668/2019-9). PLSO and HVRF are funded by a FAPESP grant (2018/00629-0). The authors would like to thank Professor Ricardo Giordano [Departamento de Bioquímica, Instituto de Química, Universidade de São Paulo (DBQ-IQUSP)] for the RGDS peptide donation and Dr Gergely Róna (Department of Biochemistry and Molecular Pharmacology, New York University School of Medicine), Dr Rebeka Tomasin (DBQ-IQUSP), Dr. Deborah Schechtman (DBQ-IQUSP), Dr. Marcelo Damário Gomes (Faculdade de Medicina de Ribeirão Preto – Universidade de São Paulo) and Dr. Hernandes F. Carvalho (UNICAMP) for all the fruitful discussions and valuable advices during the progress of this project. The authors would also like to thank Jeffrey Estrada, Celia Ludio Braga, Maria Luiza Baldini and Izaura Nobuko Toma for technical assistance.

## Author contribution

APZPF planned, performed, analyzed most experiments and co-wrote the manuscript. AMRS performed all the RT-qPCRs and immunoblottings in figure 5. PRF manufactured all hydrogels. ACM generated cDNA constructs and performed the protein stability assay. MCSB cultured the breast cell lines used in the experiments shown in figure 2A. HVRF and PSLO performed the molecular dynamics studies. MP co-supervised part of the study. AB-C planned, analyzed data, performed fluorescence microscopy, supervised the study and wrote the manuscript.

## Supplementary Figures

**Supplementary Figure 1:**
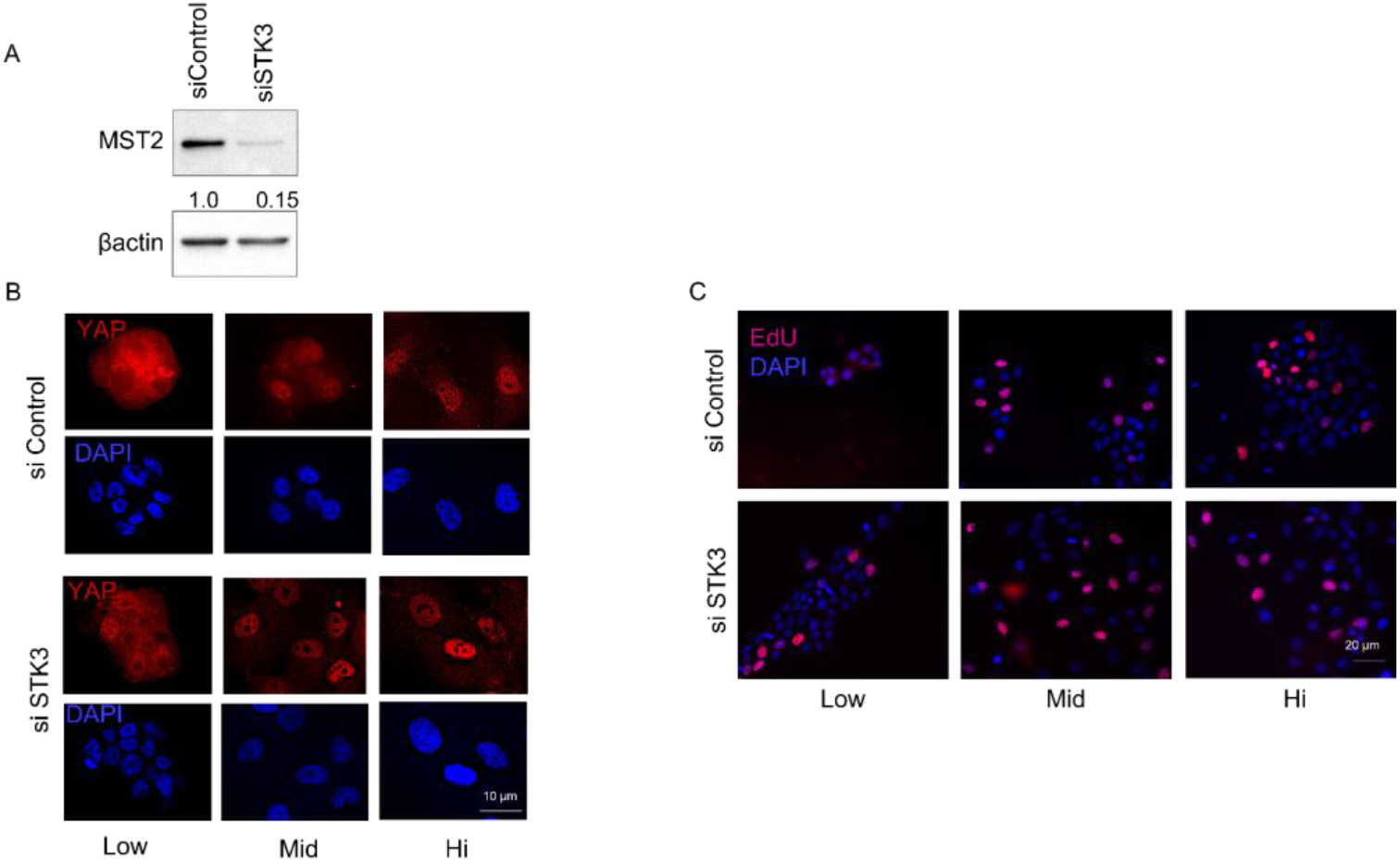
Validation of STK3 knock down in MCF10A cells and fluorescence microscopy images accompanying figure 1: A) Immunoblot for MST2 confirming STK3 silencing in MCF10A cells. MCF10A cells were cultured on regular plastic plates and treated with a control siRNA or an siRNA for STK3 for 24 hours. β-actin was used as loading control. Fold changes of MST2 are indicated under its immunoblotting. B) This panel refers to Figure 1B and displays images of YAP immunofluorescence (red) along with their corresponding nuclear staining (DAPI = blue). C) This panel refers to the quantification shown in the bar graph in Figure 1F and displays fluorescence microscopy images of EdU staining (red) along with their corresponding nuclear staining (DAPI = blue). Scale bars = B, 10 μm and C, 20 μm.

**Supplementary Figure 2:**
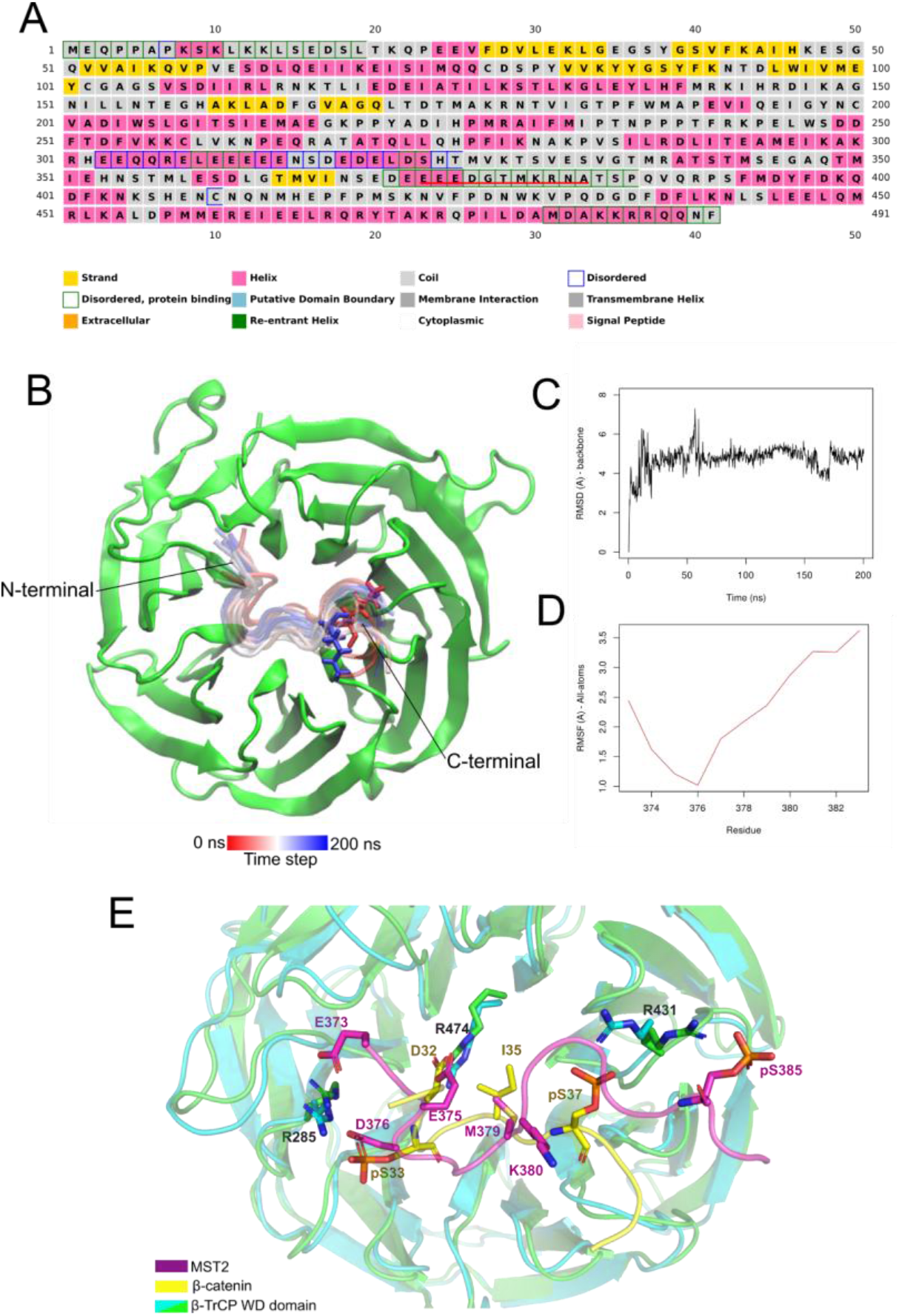
Root-mean-square deviation (RMSD) plot: A) MST2 secondary <u>structure</u> prediction. The prediction was performed in the PsiPred server. The MST2 11-mer peptide EEEDGTMKRNA is localized within a structurally disordered – protein binding region. B) 200 ns MD simulation of MST2 11-mer peptide EEEDGTMKRNA in complex with βTrCP WD40 domain. MST2 peptide snapshots of each 10 ns are shown as cartoon representation and colored by timestep. MST2 K380 is shown in sticks representing the first (red) and the last (blue) simulation frame. C) Backbone RMSD (in Å) of MST2 peptide during the MD simulation. D) RMSF calculated from all atoms of MST2 segment during the MD simulation. E) Structural alignment between β-catenin peptide crystallographic structure (PDB ID: 1P22) and MST2 peptide model bound to βTrCP. β-catenin peptide is presented in yellow and MST2 in magenta. Important residue for MST2 and βTrCP interaction are presented in sticks.

**Supplementary Figure 3:**
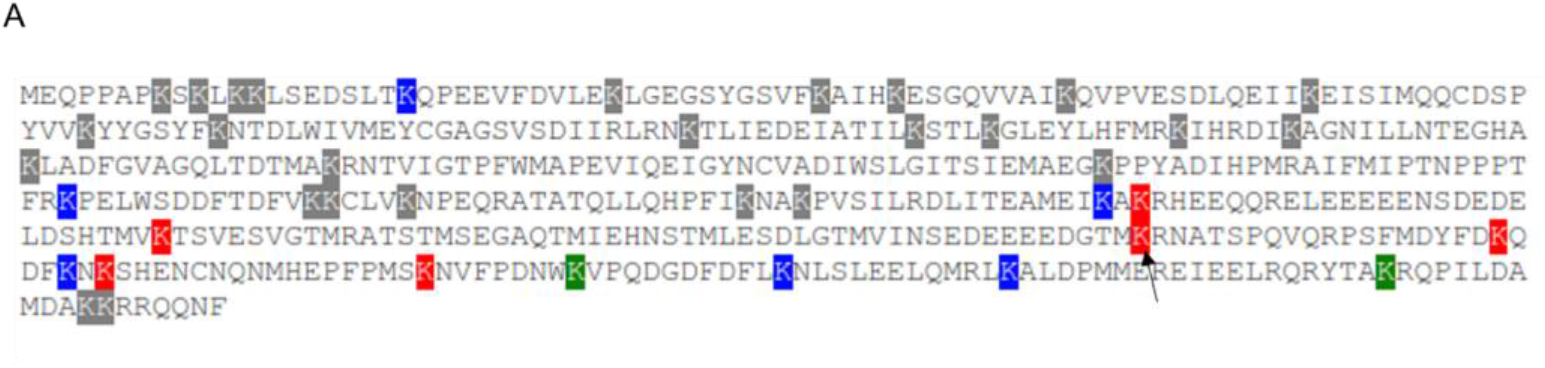
The MST2 K380 residue is predicted to be ubiquitinated: A) MST2 ubiquitination prediction. The prediction was performed using the UbPred server (http://www.ubpred.org/index.html). The colored boxes indicate the result confidence. Green, blue and red represents low, medium and high confidence, respectively. Among the MST2 lysine residues, K380 (black arrow) has the highest confidence value (0.96).

**Supplementary Figure 4.**
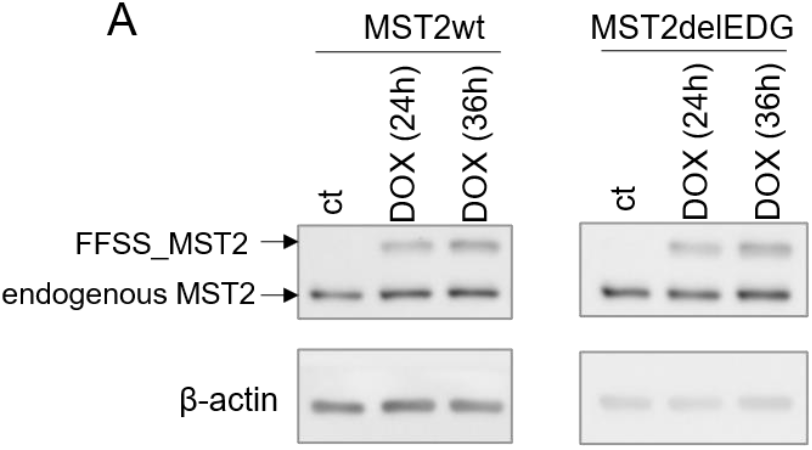
Validation of doxycycline-inducible expression of MST2wt and MST2delEDG: A) Immunoblot using an antibody for MST2 of HEK293T cells transfected with FFSS-tagged MST2wt or MST2-delEDG constructs and treated for 24 h or 36 hours with dox. Both constructs showed robust and comparable protein expression after induction.

**Supplementary Figure 5.**
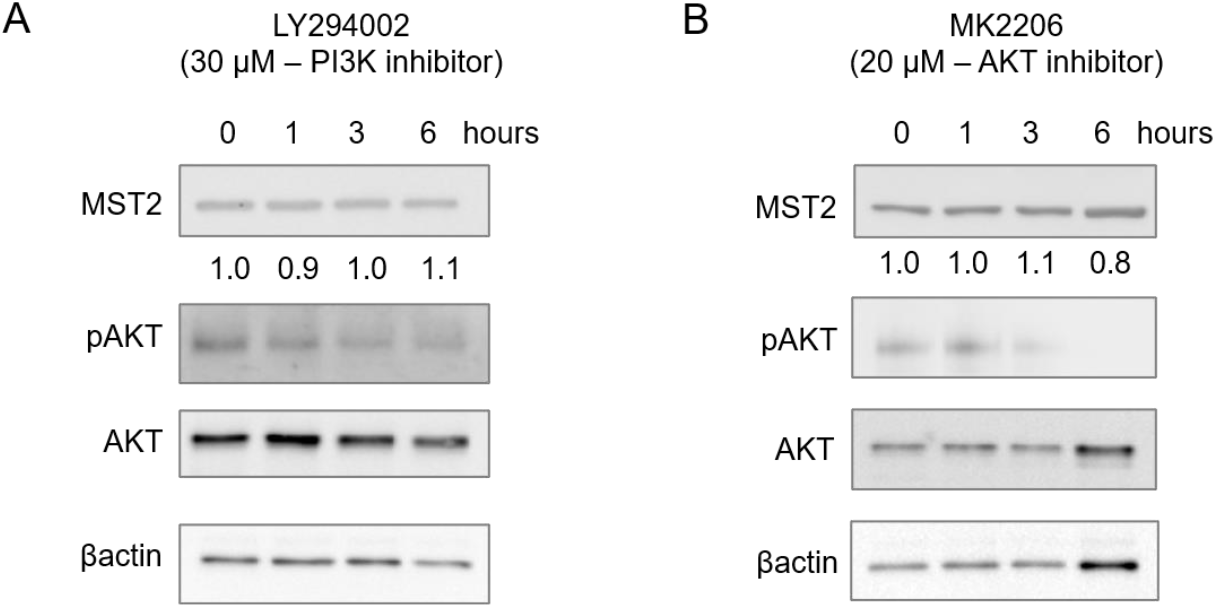
Inhibition of PI3K and AKT does not alter MST2 levels. A) Immunoblots of MCF10A cells treated with PI3 Kinase (PI3K) inhibitor (LY294002) for 1, 3 and 6 hours. AKT phosphorylation was detected to assess PI3K inactivation. β-actin was used as loading control. Fold changes of MST2 levels are shown under immunoblots for MST2 B) Immunoblot for MST2 of lysates from MCF10A cells treated with AKT inhibitor (MK2206) for 1, 3 and 6 hours. AKT phosphorylation was detected to assess its inhibition. β-actin was used as loading control. Fold changes of MST2 levels are shown under immunoblots for MST2.

## Supplementary tables

**Table 1.** List of Plasmids used in this study

| Plasmid | Backbone | cDNA cloned | Reference |
| --- | --- | --- | --- |
| pCDNA3.1_Flag_βTrCP | pCDNA3.1_Flag | βTrCP | Guardavaccaro et al 2003, (74) |
| pCDNA3.1_Flag_$\beta$ ΔFbox | pCDNA3.1_Flag | βTrCP with deletion of Fbox binding motif | Guardavaccaro et al 2003, (74) |
| pCDNA3.1_FFSS-MST2 | pCDNA3.1_Flag2xStrep2x | MST2 | This article |
| pCDNA3.1_FFSS-MST2<sup>del16-18</sup> | pCDNA3.1_Flag2xStrep2x | MST2 with deletion of EDS (amino acids 16-18) | This article |
| pCDNA3.1_FFSS-MST2<sup>del48-50</sup> | pCDNA3.1_Flag2xStrep2x | MST2 with deletion of ESG (amino acids 48-50) | This article |
| pCDNA3.1_FFSS-MST2<sup>del359-361</sup> | pCDNA3.1_Flag2xStrep2x | MST2 with deletion of ESD (amino acids 359-361) | This article |
| pCDNA3.1_FFSS-MST2<sup>del375-377</sup> | pCDNA3.1_Flag2xStrep2x | MST2 with deletion of EDG (amino acids 375-377) | This article |
| pCDNA3.1_myc_Ub | pCDNA3.1_myc | Ubiquitin | Marzio et al 2019, (66) |
| pLVX_FFSS_MST2 | pLVX-TetOne-Puro | MST2 | This article |
| pLVX_FFSS_MST2<sup>delE<sub>DG</sub></sup> | pLVX-TetOne-Puro | MST2 with the deletion of EDG (amino acids 375-377) | This article |

**Table 2.**
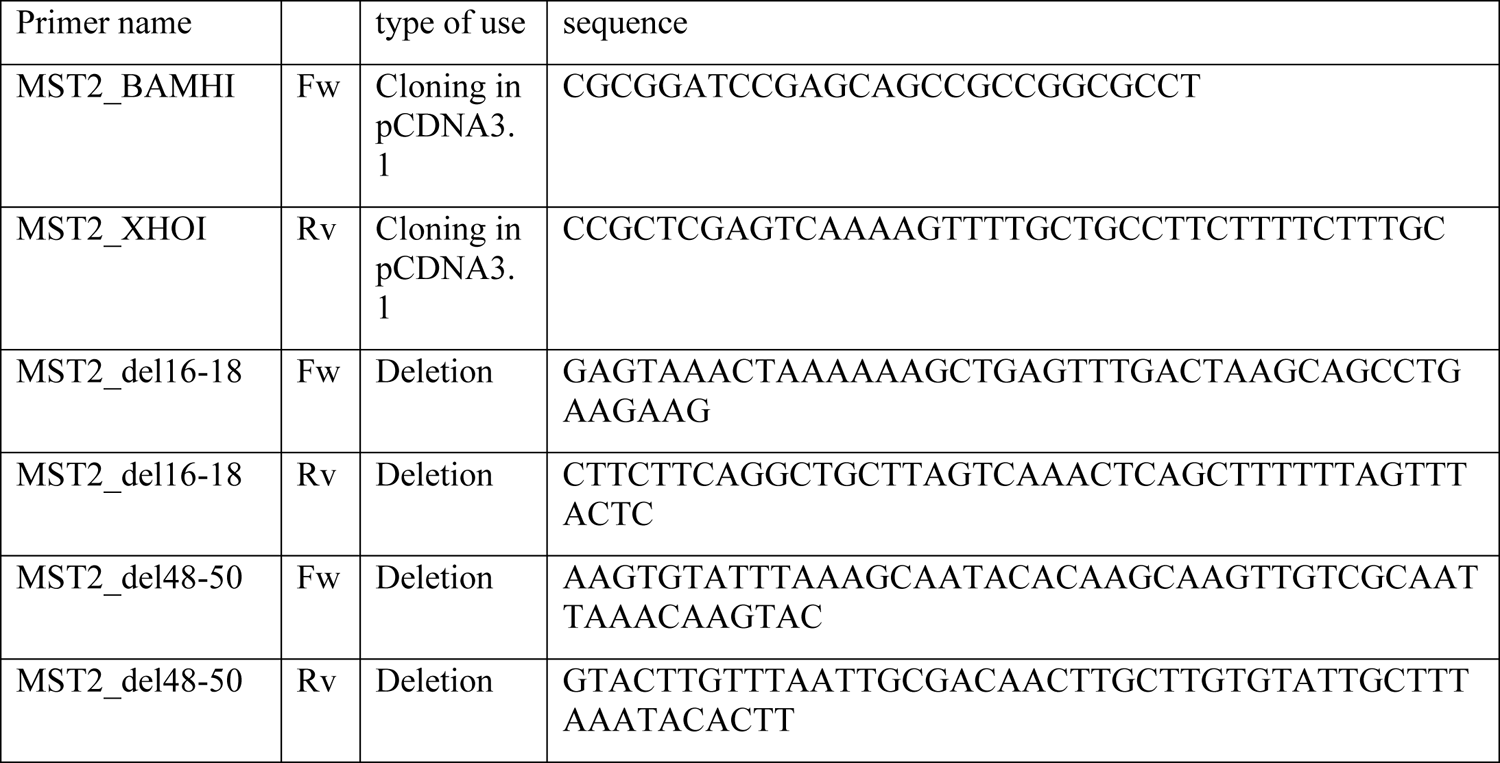
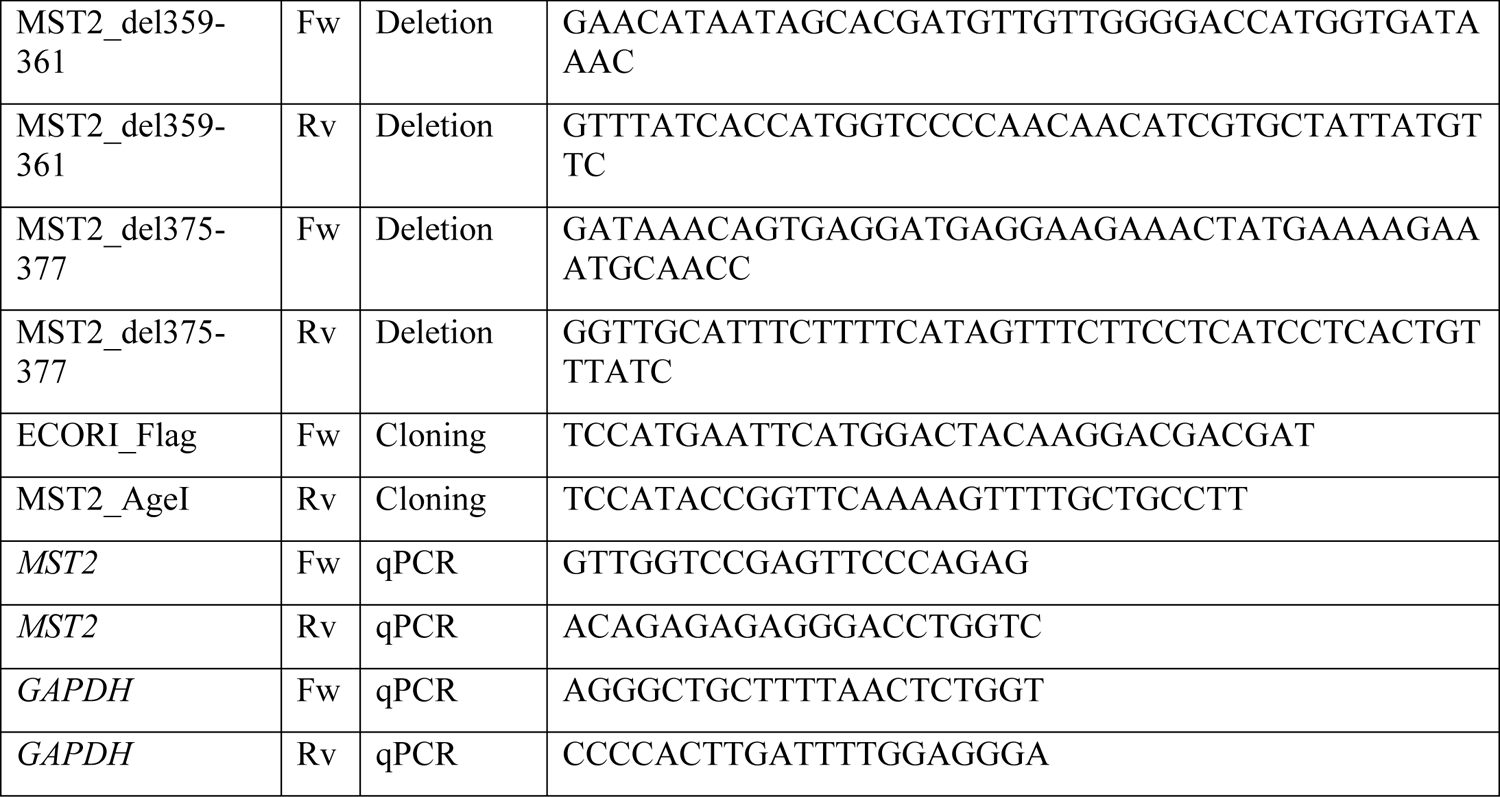
List of primers used in this study

**Table 3.**
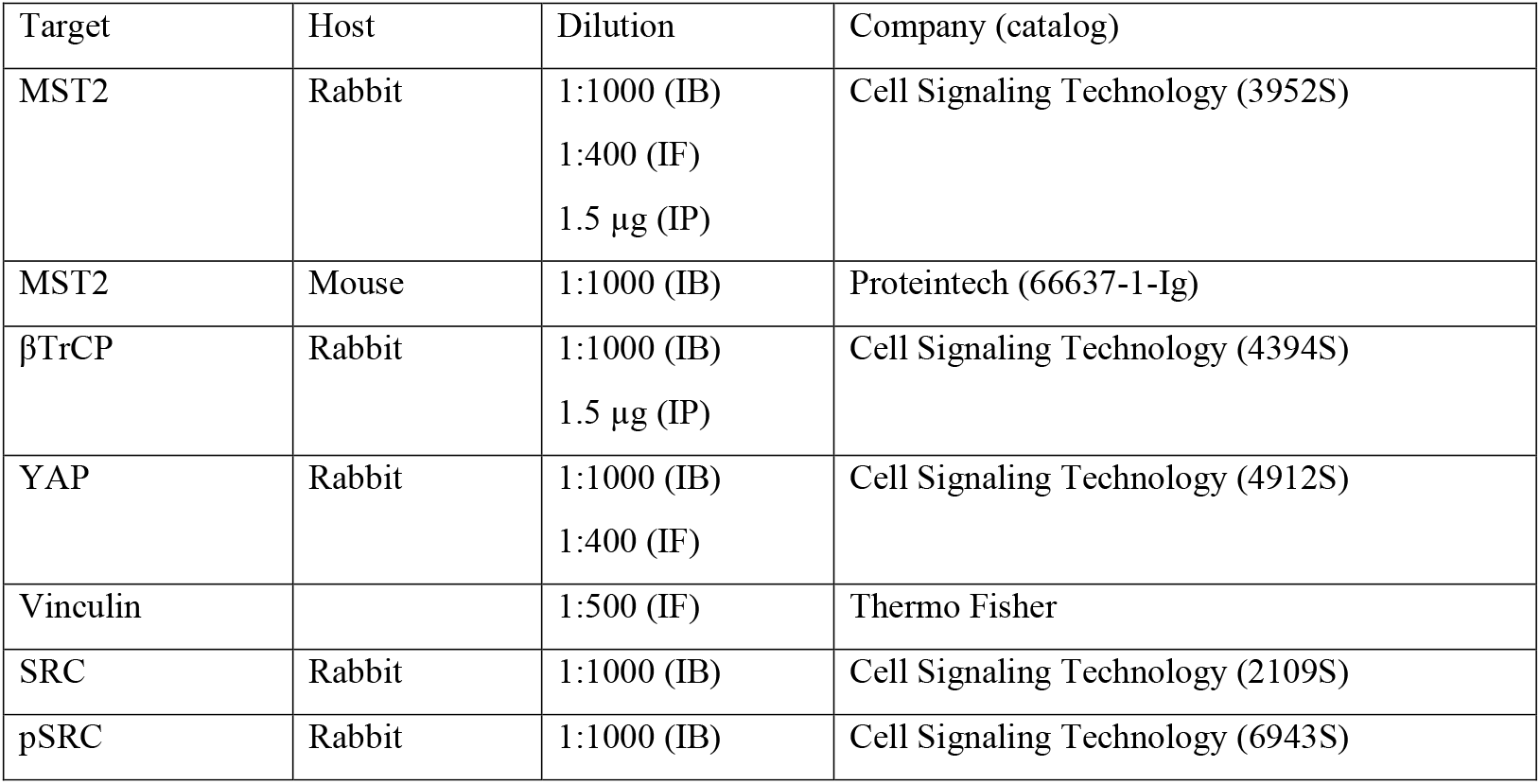
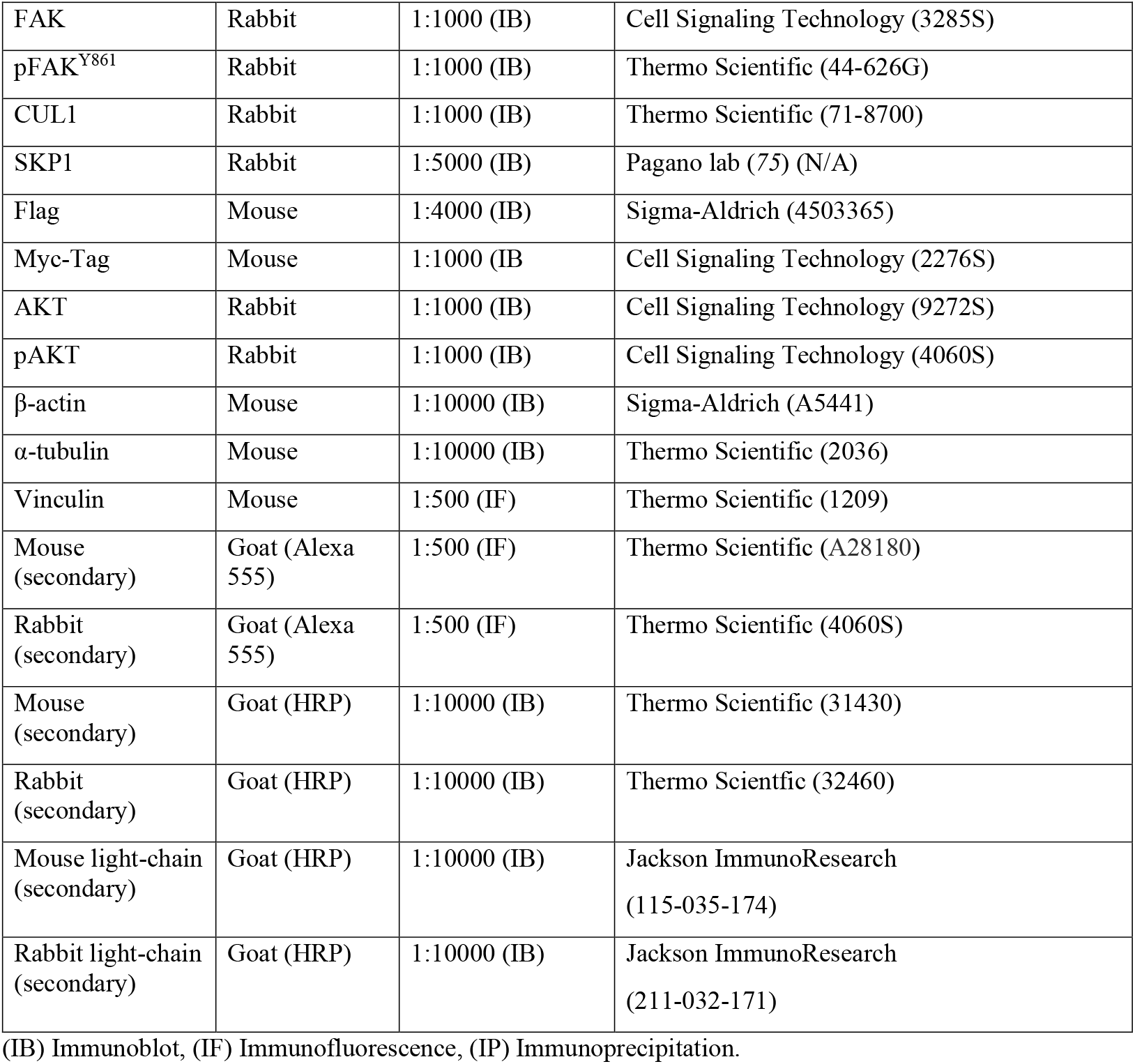
List of antibodies

| Target | Host | Dilution | Company (catalog) |
| --- | --- | --- | --- |
| MST2 | Rabbit | 1:1000 (IB)<br>1:400 (IF)<br>1.5 µg (IP) | Cell Signaling Technology (3952S) |
| MST2 | Mouse | 1:1000 (IB) | Proteintech (66637-1-Ig) |
| βTrCP | Rabbit | 1:1000 (IB)<br>1.5 µg (IP) | Cell Signaling Technology (4394S) |
| YAP | Rabbit | 1:1000 (IB)<br>1:400 (IF) | Cell Signaling Technology (4912S) |
| Vinculin |  | 1:500 (IF) | Thermo Fisher |
| SRC | Rabbit | 1:1000 (IB) | Cell Signaling Technology (2109S) |
| pSRC | Rabbit | 1:1000 (IB) | Cell Signaling Technology (6943S) |
| FAK | Rabbit | 1:1000 (IB) | Cell Signaling Technology (3285S) |
| pFAK<sup>Y861</sup> | Rabbit | 1:1000 (IB) | Thermo Scientific (44-626G) |
| CUL1 | Rabbit | 1:1000 (IB) | Thermo Scientific (71-8700) |
| SKP1 | Rabbit | 1:5000 (IB) | Pagano lab (75) (N/A) |
| Flag | Mouse | 1:4000 (IB) | Sigma-Aldrich (4503365) |
| Myc-Tag | Mouse | 1:1000 (IB) | Cell Signaling Technology (2276S) |
| AKT | Rabbit | 1:1000 (IB) | Cell Signaling Technology (9272S) |
| pAKT | Rabbit | 1:1000 (IB) | Cell Signaling Technology (4060S) |
| β-actin | Mouse | 1:10000 (IB) | Sigma-Aldrich (A5441) |
| α-tubulin | Mouse | 1:10000 (IB) | Thermo Scientific (2036) |
| Vinculin | Mouse | 1:500 (IF) | Thermo Scientific (1209) |
| Mouse (secondary) | Goat (Alexa 555) | 1:500 (IF) | Thermo Scientific (A28180) |
| Rabbit (secondary) | Goat (Alexa 555) | 1:500 (IF) | Thermo Scientific (4060S) |
| Mouse (secondary) | Goat (HRP) | 1:10000 (IB) | Thermo Scientific (31430) |
| Rabbit (secondary) | Goat (HRP) | 1:10000 (IB) | Thermo Scientific (32460) |
| Mouse light-chain (secondary) | Goat (HRP) | 1:10000 (IB) | Jackson ImmunoResearch (115-035-174) |
| Rabbit light-chain (secondary) | Goat (HRP) | 1:10000 (IB) | Jackson ImmunoResearch (211-032-171) |

(IB) Immunoblot, (IF) Immunofluorescence, (IP) Immunoprecipitation.

